# Peptide-based capture-and-release purification of extracellular vesicles and statistical algorithm enabled quality assessment

**DOI:** 10.1101/2024.02.06.578050

**Authors:** Zachary F. Greenberg, Samantha Ali, Thomas D. Schmittgen, Song Han, Steven J. Hughes, Kiley S. Graim, Mei He

## Abstract

Circulating extracellular vesicles (EVs) have gained significant attention for discovering tumor biomarkers. However, isolating EVs with well-defined homogeneous populations from complex biological samples is challenging. Different isolation methods have been found to derive different EV populations carrying different molecular contents, which confounds current investigations and hinders subsequent clinical translation. Therefore, standardizing and building a rigorous assessment of isolated EV quality associated with downstream molecular analysis is essential. To address this need, we introduce a statistical algorithm (ExoQuality Index, EQI) by integrating multiple EV characterizations (size, particle concentration, zeta potential, total protein, and RNA), enabling direct EV quality assessment and comparisons between different isolation methods. We also introduced a novel capture-release isolation approach using a pH-responsive peptide conjugated with NanoPom magnetic beads (ExCy) for simple, fast, and homogeneous EV isolation from various biological fluids. Bioinformatic analysis of next-generation sequencing (NGS) data of EV total RNAs from pancreatic cancer patient plasma samples using our novel EV isolation approach and quality index strategy illuminates how this approach improves the identification of tumor associated molecular markers. Results showed higher human mRNA coverage compared to existing isolation approaches in terms of both pancreatic cancer pathways and EV cellular component pathways using gProfiler pathway analysis. This study provides a valuable resource for researchers, establishing a workflow to prepare and analyze EV samples carefully and contributing to the advancement of reliable and rigorous EV quality assessment and clinical translation.

## 1. INTRODUCTION

Extracellular vesicles (EVs) have emerged as intriguing entities, garnering the attention of scientists from both basic and clinical research^1–6^. These small, membrane-bound vesicles hold immense potential for understanding diverse biological processes^7, 8^, such as intercellular communication, tumor initiation and progression, as well as immunity modulation. However, the isolation and purification of EVs present formidable challenges that profoundly impact downstream analysis and utility development with needed reliability and reproducibility ^9–11^, particularly for handling complex biological samples. Many biological fluids have been found to harbor an assortment of extracellular vesicle particles essential for investigations, including blood^12^, urine^13^, cerebrospinal fluid^14^, saliva^15^, pleural effusion^16^, ascites fluid^17^, amniotic fluid^18^, milk^19^, bronchoalveolar lavage fluid^20^, bacterial culture, and plant fluids. However, current EV isolation techniques are still based on sole selection property which is inefficient to precisely differentiate EV particles from other particles based on a collective outcome of size, molecular markers, and biological functionality. Furthermore, the challenges encountered in EV purification have far-reaching consequences to lead to variable and inconsistent, impairing the ability to draw accurate conclusions ^6, 9, 11, 21–33^. In the clinical sphere, the absence of standardized protocols and dependable purification methods impedes the translation of EVs for routine diagnostic and therapeutic clinical practice.

As reported by a few articles on EV isolation method comparison^9, 11, 25, 28, 34–40^, the question about which EV isolation method confers the least confounding evidence surrounding the EV associated biological effect is still unresolved, because current reports in the field have differed by sample type, metrics selected, and assay workflows, substantiating an urgent need to develop an EV isolation standardized metrics and assessment approach. Currently, the Minimum Information for the Studies in Extracellular Vesicle research guidelines^22^ (MISEV), published by the International Society of Extracellular Vesicles (ISEV) consortium, defines EV purity according to semi-quantitative metrics including particles-to-protein and particles-to-RNA relative to the reference sample, including cell media or biological sources like plasma or tissue. Additionally, assays quantifying specific EV biomarkers^22, 26, 27, 29–31, 33, 41–47^ could assist in quantifying EV purity. In consideration of non-EV contaminants, the metrics used in defining EV purity may be oversimplified as reported^9, 11, 28, 35–38^. There remains a significant gap in statistical analysis approaches that enable rigorous comparison between EV isolation methods. On the other hand, measurements in many studies are neither within the same unit scale nor consistent across preparation protocols, potentiating incomparable results that lead to confounded conclusions. Recent efforts to standardize EV isolation methods across reports have resulted in the establishment of Transparent Reporting and Centralizing Knowledge in Extracellular Vesicle Research (EV-TRACK). EV-TRACK is an approved ISEV database curating protocols from EV studies, aiming to improve comparison and reproducibility in EV research^48^; therefore, a statistical metric comparing EV isolation methods is becoming more urgently needed. We introduced an indexing strategy (ExoQuality Index, EQI) by leveraging data science to address this challenge for defining a rigorous statistical metric determined from an array of common EV isolate characteristics including nanoparticle tracking analysis (size and distribution), total particle protein, and total particle RNA, to improve the direct comparisons between isolation methods. We considered the sampling likelihood of each assay across all EV isolation methods by applying a quantile-quantile transformation^49, 50^ on the dataset, then demonstrated how EV isolation methods impact the EV quality by computing the EQI, accounting for equal likelihood of EV sampling quality across all assays and methods.

We also introduced a novel capture-release EV isolation strategy, streamlining washing steps before the on-demand release of captured EVs (ExCy) to improve EV capture specificity. We used a pH-responsive peptide conjugated on our previously developed NanoPom magnetic bead surface^41^, which crosses lipid bilayers to form an α-helix within the membrane for insertion and stabilization under acidic buffer conditions^51–56^. Different than other reported peptide EV isolation approaches^57, 58^, which use a biotin-linked irreversible insertion peptide, our EV isolation process is reversible upon restoring the buffer pH. An EV’s membrane curvature is generally smaller compared to cells. We demonstrated that EVs can be effectively captured under pH 4 with subsequent washing to remove non-EV debris. By restoring to pH 8, captured EVs can be released into a free solution. Similar acidification treatment on EV samples for improving isolation has been previously reported^59–62^, evidencing a rentention of the EV membrane fidelity at the pH used here. ExCy’s capture-release process is a simple, fast, low-cost EV isolation workflow requiring only a magnet, acidic buffer and dye, centrifuge, and pipette.

We employed ExCy to purify a variety of biological fluids, including human patient plasma, mammalian cell culture medium, cow milk, bacterial culture for outer membrane vesicles, orange juice, and hemp juice to demonstrate broad applicability (Figures 1-2 and Figure s1). To demonstrate the EQI’s cross-platform capability for evaluating isolation quality between isolation methods, we applied the EQI to compare pancreatic cancer patient plasma and healthy control isolated EVs from ExCy with those EVs isolated from membrane affinity (ExoEasy Maxi Kit), sucrose cushion ultracentrifugation (UC), and phosphatidylserine affinity bead capture (Fujifilm MagCapture). Furthermore, we conducted total EV RNA sequencing and transcriptomic analysis to compare the landscape of human plasma EV-derived RNAs across all methods, as well as the transcriptomic analysis of tumor biomarkers from pancreatic cancer patients with healthy individuals as the control. We identified ATP6V0B gene as uniquely expressed from pancreatic patient plasma EVs using our isolation method when compared to other three isolation methods, which is a well recognized oncogene mediating eukaryotic intracellular organelles, including protein sorting, zymogen activation, receptor-mediated endocytosis, and synaptic vesicle proton gradient generation (Human Protein Atlas)^63^. Therefore, our results support the higher specificity and coverage range of our developed EV isolation method (ExCy) in terms of alignment with pancreatic tumor pathways and EV cellular component pathways.

**Figure 1.**
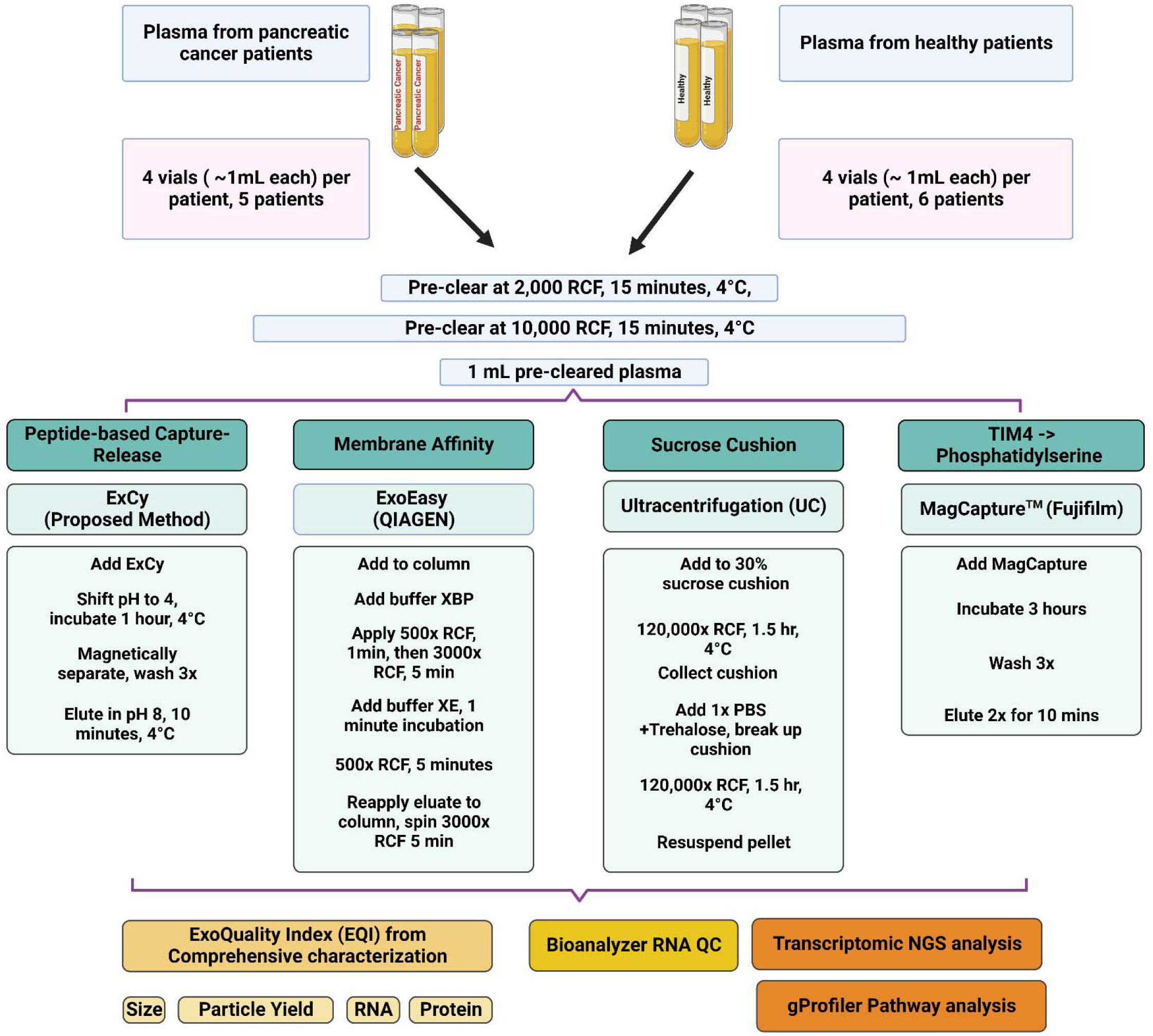
Workflow of sample preparation of EVs from human patient plasma associated with the quality index assessment and transcriptomic NGS study.

**Figure 2.**
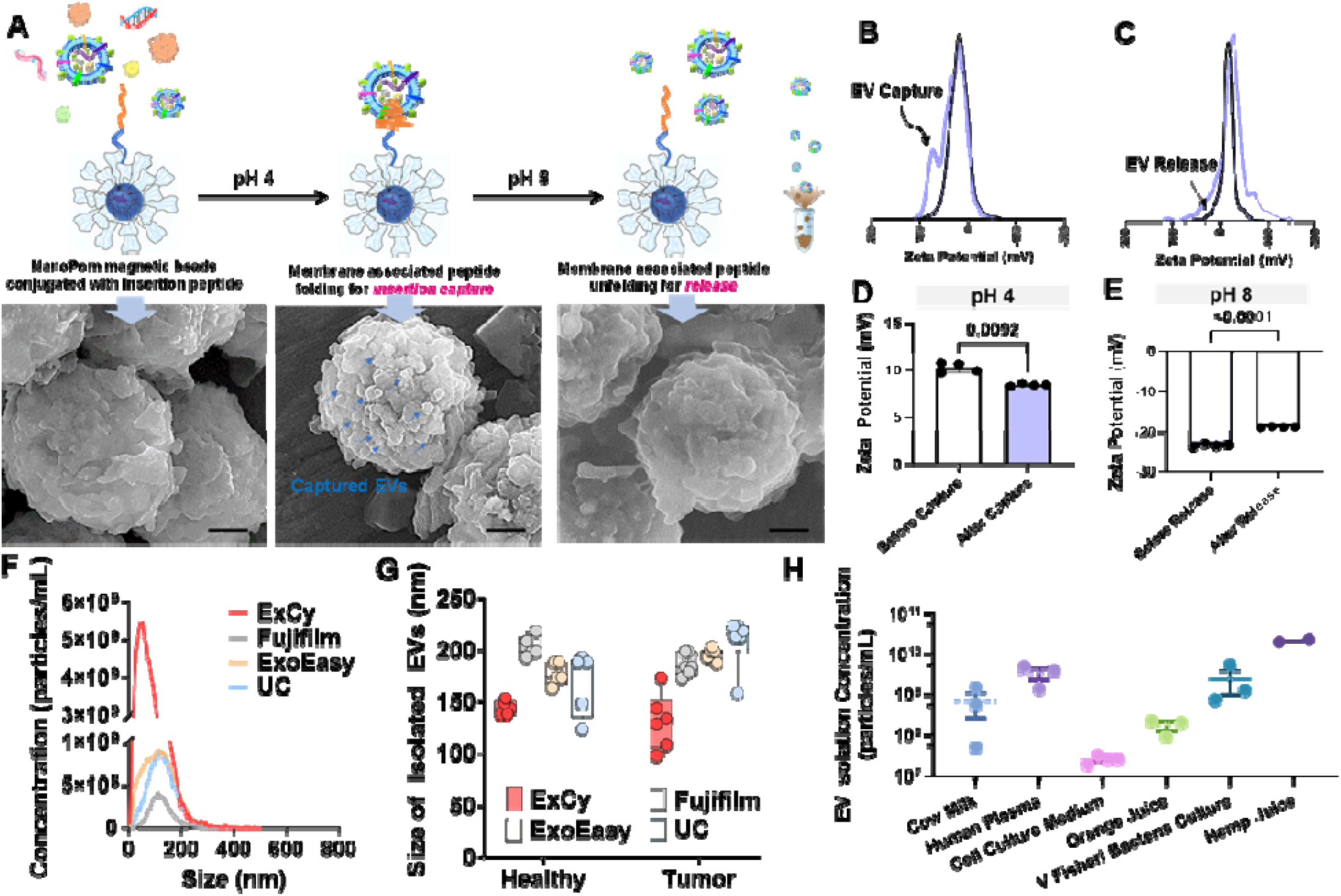
ExCy’s proof-of-principle of capturing and releasing EVs. **A**. Schematic illustration of ExCy’s reversible capture-and-release of EVs with demonstration via SEM imaging. **B**. ExCy’s zeta potential distribution profile during the capture and **C)** release steps. **D**. ExCy’s zeta potential before and after EV capture. **E**. ExCy’s zeta potential before and after EV release. **F**. Nanoparticle tracking analysis of isolated EV concentration compared between ExCy, Qiagen’s ExoEasy, Kodak’s Fujifilm, and ultracentrifugation (UC). The pancreatic cancer patient plasma samples were used. **G**. The size of EVs isolated from pancreatic cancer patient plasma samples compared across all methods (n=6 for tumor and 5 for healthy, mean ± sd), indicating ExCy captures more small sized plasma EVs below 150-200 nm than ExoEasy, Fujifilm, and UC. **H**. ExCy isolation of EVs from a variety of biological fluids to prove the reproducibility and applicability of developed isolation approach.

## 2. MATERIAL AND METHODS

We have submitted all relevant data of our experiments to EV-TRACK (EV-TRACK ID: EV240005)

### 2.1. Plasma Collection & Sample Preparation

The Clinical and Translational Science Institute, University of Florida, Gainesville, FL, collected blood plasma from pancreatic cancer patients and healthy individuals under the approved IRB protocol (IRB#202102401) and patient consent. The samples were thawed from –80°C for processing. Sample processing and characterization were performed on the same day to avoid environmental change-associated variations. As illustrated in Figure 1, two rounds of pre-clearing low-speed centrifugation were applied to remove cellular debris. Final volumes for all pre-cleared plasma samples were 1 mL.

### 2.2 ExCy Bead Fabrication and Isolation

The pH-responsive peptide conjugated beads comprise our previously published unique nanographene pom pom-like microbeads (NanoPom)^41^. Briefly, the NanoPom is fabricated by using a Fe_3_O_4_/SiO_2_ core, followed by a series of sequential layer depositions of nanographene to form pom-like sheet layers with nanocavity in between, which enables a size hindrance to avoid considerably larger membrane structures. Streptavidin is cross-linked onto the beads, allowing a biotinylated insertion peptide conjugation to form ExCy beads. The pH-responsive peptide was synthesized through the KU Molecular Probe Core using a peptide microwave synthesizer and purified and analyzed by LC-MS. The quality and yield of synthesized peptides were characterized in Figure s2.

To capture EVs, 40 μl (approx. 0.4 mg beads) of ExCy was mixed with 1 mL of plasma. Depending on the original pH of biological sample fluids, the corresponding volume of HCl in 3M concentration was added to the sample solution along with 5 uL of pH indicator Methyl Purple to achieve a buffer pH of around ∼4. Detailed validation for various biological fluids was illustrated in Figure s1. Afterward, the sample mixture was incubated on a revolver for 1 hour at 4°C. Methyl purple was added to ensure the pH of the buffer between 4-6. After incubation, captured EVs were magnetically aggregated to remove the liquid solution for washing 3 times with ice-cold, pH 4, 1x PBS. Subsequently, the washed beads solution in 40 μl was transferred to a 200 uL solution of pH 8, 1x PBS containing 10 mM Tris for gentle vortex and release to harvest intact EVs. The detailed protocols, along with specific biological sample types, were shown in Figure s1.

### 2.3 ExoEasy Maxi Kit for EV isolation

ExoEasy (QIAGEN) was used according to the manufacturer’s protocol. 1 mL of plasma was added to an equal volume of XBP reagent, mixed, and centrifuged at 500x RCF for 5 minutes at room temperature. The membrane was washed with XWP reagent using 3000x RCF for 5 minutes at room temperature, followed by adding 400 uL of XE reagent for a 5-minute centrifugation at 500x RCF at room temperature. We reapplied the eluate to the column, as per manufacturer recommendation, and centrifuged for 3000x RCF for 5 minutes to obtain EVs.

### 2.4 Fujifilm MagCapture for EV isolation

Fujifilm’s MagCapture Exosome Isolation Kit PS v1 was used per the manufacturer’s protocol. 60 uL of magnetic beads were combined with 500 uL of Exosome Capture Immobilizing Buffer, mixed, and supernatant removed. 10 uL of Biotin-labeled Exosome Capture reagent was added to the beads in 500 uL of Exosome Capture Immobilizing Buffer for 10 minutes at 4°C. The beads were washed 3x with 500 uL of the Exosome Capture Immobilizing Buffer. Notably, 2 uL of Exosome Binding Enhancer was added to 1 mL of plasma sample. Washed beads were magnetically removed, applied to the plasma sample, and incubated over 3 hours at 4°C. After incubation, 6 uL Exosome Binding Enhancer was added to 3 mL of Exosome Washing Buffer. This washing buffer was used to wash EV-captured beads 3x, followed by eluting twice with 50 uL of the Exosome Elution Buffer for a final eluate volume of EVs in 100 uL.

### 2.5 Sucrose Cushion Ultracentrifugation (UC)^64, 65^

A 3 mL 30% (w/v) sucrose solution was placed at the bottom of an ultracentrifuge tube containing 15 mL 1x ice-cold PBS. Next, 1 mL of plasma was gently added to the ultracentrifuge tube to not disturb the sucrose cushion, followed by a round of ultracentrifugation at 120,000x RCF for 1.5 hours at 4°C. Next, the sucrose cushion was transferred to a different ultracentrifuge tube, 15 mL of 1x ice cold PBS was added, followed by pipetting to gently break the sucrose cushion and ultracentrifuged again at 120,000x RCF for 1.5 hours at 4°C. The supernatant was removed, followed by pellet resuspension in 200 uL 1x PBS to collect EVs.

### 2.6. Preparation for Downstream Analysis

After EV purification using four different isolation methods (ExCy, ExoEasy, FujiFilm, UC) from the same patient samples, all isolates received 25 mM PBS-trehalose ^23, 66, 67^ up to a final volume of 500 uL. Next, 300 uL was used for NTA, 18 uL for protein extraction, 5 uL for TEM, and 77 uL for RNA extraction. Each measurement was conducted 4 independent times with 4 replicates per measurement.

### 2.7 Scanning Electron Microscopy

Field emission-scanning electron microscopy (FE-SEM) was used for visualizing the bead morphology, topology, and EV sizes, pre– and post-EV capture. Pre-capture beads were washed 3x with ice-cold pH 4, 1x PBS, and resuspended in 25 uL washing buffer. Post-capture beads were washed 3x in ice-cold pH 8.0 1x PBS with 10 mM Tris and resuspended in 25 uL of releasing washing buffer. Each resuspension solution was gently mixed and then directly aliquoted onto a 100% acetone-cleaned Ted Pella aluminum pin stub mount with complete solution evaporation. Next, the samples were sputter-coated for 30 s with an Au/Pd target using a Denton Desk V Sputter Coater and loaded into a Hitachi SU5000 Schottky Field-Emission Scanning Electron Microscope at a high negative vacuum pressure of 10-8 torr. An incident electron beam was applied to the samples at 7 keV and a beam current of 16.7 nA. Aperture and stigmata corrections were done before sample images were obtained.

### 2.8 Transmission Electron Microscopy

Transmission electron microscopy (TEM, FEI Spirit TEM 120 kV) verified the morphology of isolated EVs. Briefly, ultrathin copper grids coated with 400 mesh carbon film (FCF400-Cu-UB, Electron Microscopy Science, USA) were used with glow discharge treatment for 1 min before use. Then, 5 μl EV samples were individually added onto the glow-discharged grids and were quiescent for 10 min at room temperature. The grids were washed with distilled water once, then negatively stained with filtered 2% aqueous uranyl acetate for 2 min, and dried at room temperature before observation. The TEM imaging power was 120 kV by FEI Spirit G2 with a digital camera (Soft Image System, Morada and Gatan Orius SC 1000B CCD-camera).

### 2.9 Nanoparticle Tracking Analysis

Nanoparticle tracking analysis was conducted using ZetaView (QUATT, Particle Metrix Inc, USA). ZetaView measures the nanoparticle’s Brownian motion using an incident laser to determine the corresponding size. The nanoparticle motion is then tracked by the detector and recorded over time: The incident laser wavelength was 488 nm^-1^ with sensitivity at 75 and shutter time at 163, over 90 seconds at the highest video resolution for all 11 positions.

### 2.10 Zeta Potential Measurement

Zeta potential was measured using the Litesizer 500 (Anton Paar, Austria). The Litesizer 500 measures the zeta potential by flowing an electric current between two electrodes within the cuvette, measuring the interfacial charge of the solution at the particle’s surface and correlating this to the particle’s surface charge. ExCy beads under different conditions were dispersed within a 0.1x PBS solution at pH 4 or 8 for measurement.

### 2.11 Protein Extraction and SDS-PAGE

18 uL of Isolated EVs were lysed with 2 uL of 1x ice-cold RIPA buffer. Samples were incubated on ice for 15 minutes, with 30-second vortexing every 5 minutes. At the end of the incubation, the samples were sonicated for 15 seconds. Total protein was then quantified with the Pierce BCA Protein Assay Kit (Thermo Fisher, USA). Per the vendor’s protocol, reagents A and B were mixed at a 24:1 ratio to formulate the working buffer. In a separate tube, 10 uL of the working reagent and lysed EV sample were added, followed by incubation at 75°C for 5 minutes, then cooled to room temperature to measure 562 nm^-1^ absorbance. Known quantities of bovine serum albumin were utilized to generate the standard calibration curve. For SDS-PAGE, 1 uL of Halt’s 100x Protease Inhibitor Cocktail was supplemented to lysed samples. Approximately 20 ug of protein from each sample was resolved on a 4-20% gradient gel (Biotek), then visualized by SimplyBlue SafeStain.

### 2.12 RNA Extraction & Bioanalyzer

Total RNA was extracted using the miRNeasy Mini Kit per manufacturer instructions from EVs pre-processed with 5 uL of 1x DNase I and RNase buffer. Then, isolated EVs were lysed with QIAzol and incubated for 5 minutes at room temperature. Next, chloroform was added to separate the lipids, mixed vigorously, then incubated at room temperature for 3 minutes, followed by 12000x RCF centrifugation for 15 min at 4° to allow phase separation. The aqueous phase was extracted into a separate tube, mixed with 100% ethanol, and then transferred to the RNeasy Mini column. The column was centrifuged at 8000x RCF for 15 seconds at room temperature, followed by washing with 700 uL of reagent RWT and centrifuging at 8000x RCF for 15 seconds at room temperature. Next, 500 uL of reagent RPE was added to the RNeasy column and centrifuged for 2 minutes at 8000x RCF. We further dried the membrane by centrifuging an additional time at full speed for 1 minute. Afterward, 50 uL of RNase-free water was added to the column to elute the RNA from the membrane using 8000x RCF for 1 minute. The eluate was applied to the membrane an additional time. 5 uL of 1x DNase I was added to remove any leftover DNA, followed by measuring the RNA quality through the 260/280 nm^-1^ ratio (BioTek Cytation 5). Each sample’s RNA was then analyzed using Agilent’s 2100 Bioanalyzer (Agilent Technologies) with the total RNA 6000 Pico Kit, according to the manufacturer’s protocol.

### 2.13 Next Generation Sequencing – NovaSeq 6000

All extracted RNA samples, analyzed from the Bioanalyzer, for the inputs to NEBNext® Ultra RNA Library Prep Kit, were prepared as 2 ng / uL concentrations for cDNA preparation and adaptor ligation per manufacturer instructions. Briefly, 50 uL of sample was used to generate the first cDNA strand with the thermocycler settings of 25°C, 42°C, 70°C, and 4°C for 10, 15 x2 minutes, and then held, respectively. The second cDNA reaction was immediately performed at 16°C for 1 hour with the heated lit at 40°C. NEB’s SPIR magnetic beads purified double-stranded cDNA, then combined with the End Prep Reaction mix and enzyme, mixed, and incubated in a thermocycler at 20°C, 65°C, and 70°C for 30 x2 mins, and held until the adapter ligation reaction. Adapter, ligation master mix, and enzymes were mixed with the samples on ice, followed by incubation in the thermocycler at 20°C for 15 minutes. The USER enzyme was added to the samples, mixed, and incubated at 37°C for 15 minutes with the heated lit set to 45°C. Samples were purified with SPIR magnetic beads again. Adapter ligated DNA was subjected to PCR after adding the NEBNext® Multiplex Oligo 96 Unique Dual Index Primer Pairs, adding unique i7 and i5 forward and reverse primers to each sample. The Qubit Fluorometer quantified the constructed RNA library, and the Bioanalyzer checked the quality. Finally, all samples with the unique bar codes were pooled and sequenced by Nova Seq 6000.

### 2.14 Transcriptomic Analysis

Transcriptomic analysis was conducted through the FREYA pipeline^68^. Briefly, reads were mapped to the hg38 genome with HISAT2^69^, followed by assembly quality control using FastQC, Trimmomatic^70^, and GATK tools^71^ Picard and SplitNCigarReads. Next, DEXSeq-Count^72, 73^ was used to generate the read counts for each transcript. Transcripts less than an average of 10 reads were filtered from the analysis. Differential expression testing was conducted by edgeR^74–76^, which uses a negative binomial distribution for the generalized linear model explaining EV isolate differences while adjusting for each method’s contribution to the observed EV isolate’s mRNA. Within edgeR, samples were normalized using the weighted trimmed mean of M-values^77^ and the false discovery rate (<0.05) for statistically different transcripts was calculated by Benjamini-Hochberg^78^. Transcripts passing the false discovery rate cutoff were used in downstream enrichment analysis. Enrichment analysis of statistically significant transcripts was performed using gProfiler^79^ and the HumanBase^80^ community detection algorithm to identify tissue-specific functional network interactions enriched with differentially expressed transcripts. HumanBase builds genome-scale functional map of human tissues to serve as a profiler of gene-specific function within tissue networks. HumanBase will attempt to profile input genes with any interacting partners, illustrated as the interaction confidence.

### 2.15 Statistics

A two-tailed *t* test was conducted to determine ExCy’s capture of EVs by zeta potential in Figure 1. All plots with error bars mean ± standard deviation (N=4 per sample). Analyses and plotting were performed with Python v3.8, R v4.1 and GraphPad v9.0.

## 3. RESULTS

### 3.1 The pH responsive peptide enabled capture-release NanoPom magnetic beads (ExCy) for purifying EVs

ExCy magnetic beads are building upon our previously reported NanoPom^41^ magnetic beads by introducing the conjugation with pH-responsive peptides, which confers a high surface area for capture and nanocavities favorable to nano-sized EVs, in turn, for isolating a well-defined EV population. The pH-responsive eptides have spontaneous translocation ability to cross the lipid bilayer and form an α-helix in the membrane for insertion and stabilization under acidic buffer conditions^51–56, 81^, while alkalizing to release in pH 8. As illustrated in Figure 2A and validated using SEM, the EV capture and release can be visualized on the bead surface before and after buffer pH changes. Since ExCy-captured EVs can undergo multiple rounds of washing to remove protein aggregates and cellular debris before EV release, the EV population homogeneity remains high. We also measured the zeta potential profiles and associated changes before and after the capture or release of EVs from ExCy beads, as shown in Figure 2B-E for validation, which showed distinctive profiles and significant changes indicating the event from EV attachment. The zeta potential change could support the surface property change due to the event of EV attachment, which leads to a second peak in the zeta otential distribution profile due to the expanded Debye length by EV attachment (Figure 2B) rather than a singular broad peak often observed with plasma protein debris in colloidal systems^82–84^. On the other hand, we also observed a significant decrease in peak width (Figure 2C) for the event of EV release. The zeta potential validation is consistent with our SEM observations. The nanoparticle tracking analysis of isolated EVs showed higher abundant particle concentration and comparable size range compared to conventional isolation methods, as detailed in Figures 2F and G. In addition to isolation from pancreatic cancer patient plasma samples, we also tested a variety of biological fluids, including human patient urines, mammalian cell culture medium, cow milk, bacterial culture for outer membrane vesicles, orange juice, and hemp juice, as summarized in Figure 2H and Figure s1, showing broad applicability. The reusability of these ExCy capture-release beads was also assessed to show consistent EV isolation performance after four cycles of capture-release, as demonstrated in Figure s3.

### 3.2 Statistical algorithm (ExoQuality Index, EQI) for assessing EV isolation quality

EV research regarding isolation and analysis is often subject to significant variation. Different isolation methods apply different preparation protocols. Furthermore, different assays measuring an EV’s protein, RNA, and yield with different preparation protocols and instruments could induce chained differences, which ar highly significant to majorly confound the investigation’s conclusions. Thus, standardization and cross-comparison are essential in the field. To address this gap, we introduced a direct comparison metric for assessing EV isolation quality by employing a statistical algorithm and creating an indexing strategy that intakes MISEV-suggested measurements^85^, including total RNAs, proteins, yield, and size distribution (Fig re 3A, Supplementary File 1). First, we applied each isolation method on purifying 11 human plasma samples (6 pancreatic cancer patient plasma samples and 5 healthy plasma controls) to tabulate total RNAs (Figure 3B), proteins (Figure 3C), and particle yield (Figure 3D), size and size distribution for analysis (Figure s4). The quality of isolated total EV RNAs and proteins from different isolation methods was further assessed by 260/280 nm^-1^ absorbance ratio (Figure s5) and SDS-PAGE gel electrophoresis (Figure s6), which indicated the high-quality yield using our developed ExCy isolation method.

**Figure 3.**
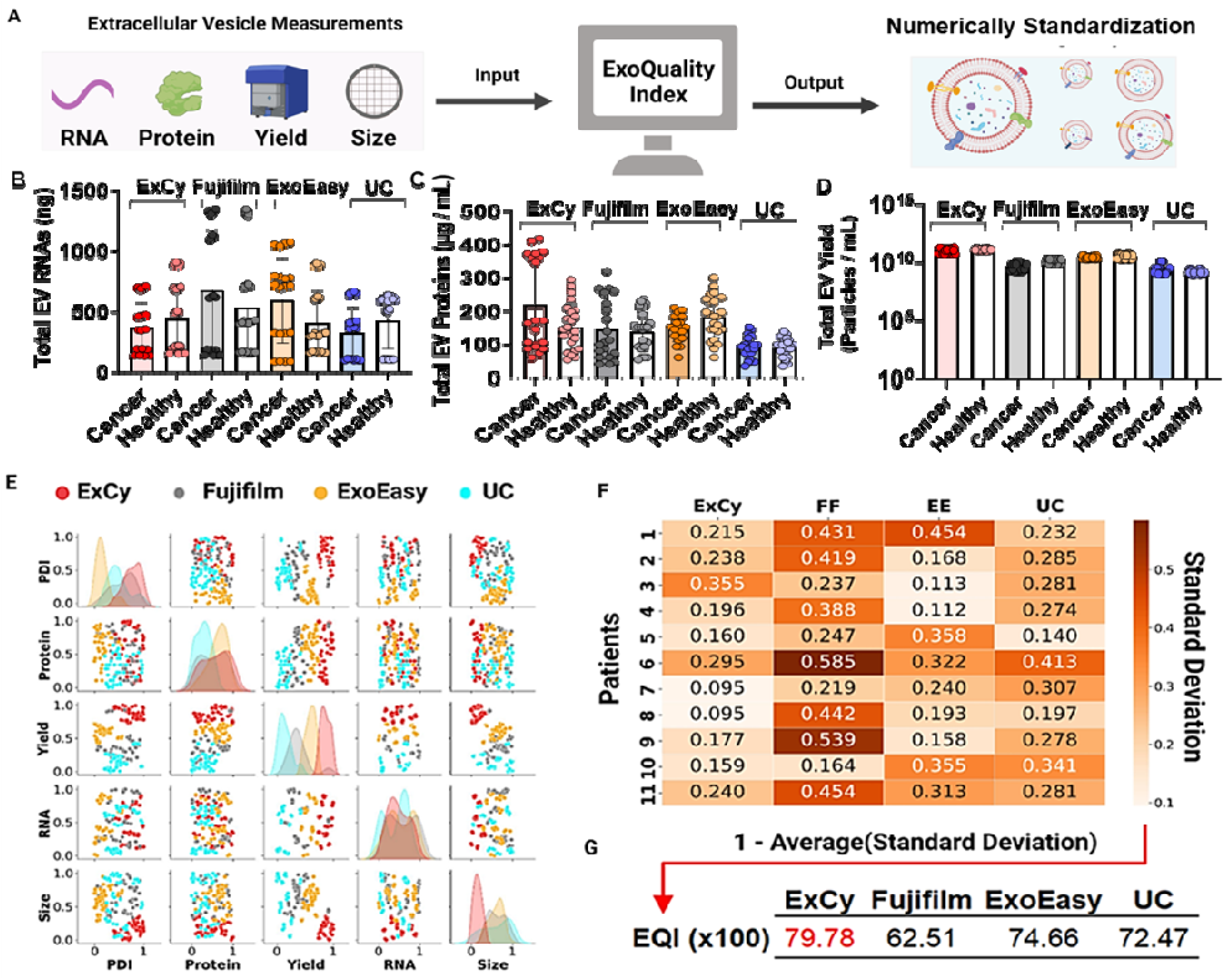
Statistical analysis using ExoQuality algorithm (EQI) to standardize EV isolation method for comparison. **A**. Schematic illustration on how ExoQuality to evaluate quality of EV isolates based on EV characterizations. **B**. Stratification of patients to further investigate method-dependent observed total EV RNAs, **C)** EV proteins, and **D)** EV particle yield. **E**. Scatter plot matrix comparing the association of each EV measurement across the isolation methods. The particle dispersity index (PDI)^86^ is a representation of the EV size distribution within a sample population, detailing observed heterogeneity. PDI was calculated as 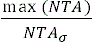 where NTA is the size profile and σ is the standard deviation. All isolations were done at the same date and location. The data was then transformed to a uniform distribution to adjust for each method’s bias contribution, applying a probabilistic rescaling within each EV measurement. **F**. A sample-by-method standard deviation matrix was calculated from each method’s covariation matrix per sample. Within the probalistic rescaling, a standard deviation value of 0.5 represents a likelihood to observe 50% deviation after resampling. **G**. The ExoQuality index metric used for direct comparison between EV isolation methods, intepreted as the expected likelihood of EV homogeneity, after resampling to observe similar measurements.

Generally, the size profiling and protein and RNA quantitation measurements are in different units and not on the same measurement scale. Therefore, we applied a quantile transformation to the data to standardize both units and measurement scales (Figure 3E). A quantile transformation transforms all the data based on the cumulative distribution function, enabling direct comparisons across measurements through a probabilistic perspective ^49, 50^. As seen by reported EV isolation comparison articles ^9, 11, 25, 28, 34–40^, we also observed distinct EV populations from different isolation methods. For example, distinct EV populations relative to their isolation methods were observed when visualizing total RNAs or proteins against EV isolate yield, size, and size distribution. Additionally, Figure 3E’s diagonal represents the univariate distribution of each EV quality measurement within an equal likelihood space, indicating how each measurement could be distinctive depending on the isolation method. From this analysis, EV total RNAs and proteins are two nearly independent metrics on the isolation methods. However, an EV’s size, yield, and size distribution would suggest method-dependent observance on isolated EVs, which could be due to the semi-quantitative nature of NTA measurements, which may confound the study conclusions.

To address the incongruencies on analyzing an isolated EV population when given multiple assays and isolation methods, we computed the sample-by-method standard deviation matrix (SMSD) after quantile transformation to attempt standardization between EV isolation methods and enable direct comparison. The SMSD matrix was calculated by computing the covariance matrix for each method per patient, assuming the parametric variance followed by square rooting and tabulating (Figure 3F). Our analysis illustrated each isolation method’s impact on EV isolate quality. For example, in our dataset, a value of 0.2 indicates that the expected EV population may differ by 20% after resampling with the isolation method. For instance, in a pairwise sample comparison, Fujifilm deviates more frequently than ExCy, ExoEasy, and UC. Therefore, we further calculated a method’s expected EV isolate standard deviation for interpretation as the *ExoQuality Index = 1 – Average(Standard Deviation)*, attempting to directly quantify and compare between methods (Figure 3G). The result indicates ExCy (EQI 79.78%) has the most homogeneity of isolated EVs with least deviation after resampling, suggesting the most consistent population from an EV isolation.

### 3.3. NGS for exploratory analysis of EV isolation quality

After performing the EQI index analysis, we also used NGS to investigate the transcriptomic profiles from different EV isolates (Figure 4). We first selected early-stage pancreatic patient plasma samples (n=4) with healthy plasma (n=3) as the control to isolate EVs using 4 different isolation methods with subsequent extraction of associated total RNAs, respectively. The quantification of a total of 28 RNA extracts using the Bioanalyzer was shown in Figures s7-10. Utilizing the FREYA pipeline^68^, we investigated the raw sequence assembly quality through FastQC (Supplementary File 2), followed by mapping to hg38 using HISAT2^69^. HISAT2’s mapping rate across the methods indicated ExCy, on average, had the highest hg38 alignment rate and the least varied mapping rate (Figure s11), suggesting purer EV isolated RNA quality. Following, we then tabulated the RNA population in each patient across all methods (Figure s12). We further quantified method-dependent EV RNA annotations by querying to the MISEV-approved EV database, Vesiclepedia^87^ to understand levels of non-EV reported annotations (Figure s13) and further increase the analytical rigor in the following analysis by standardizing method-dependent EV RNAs.

**Figure 4.**
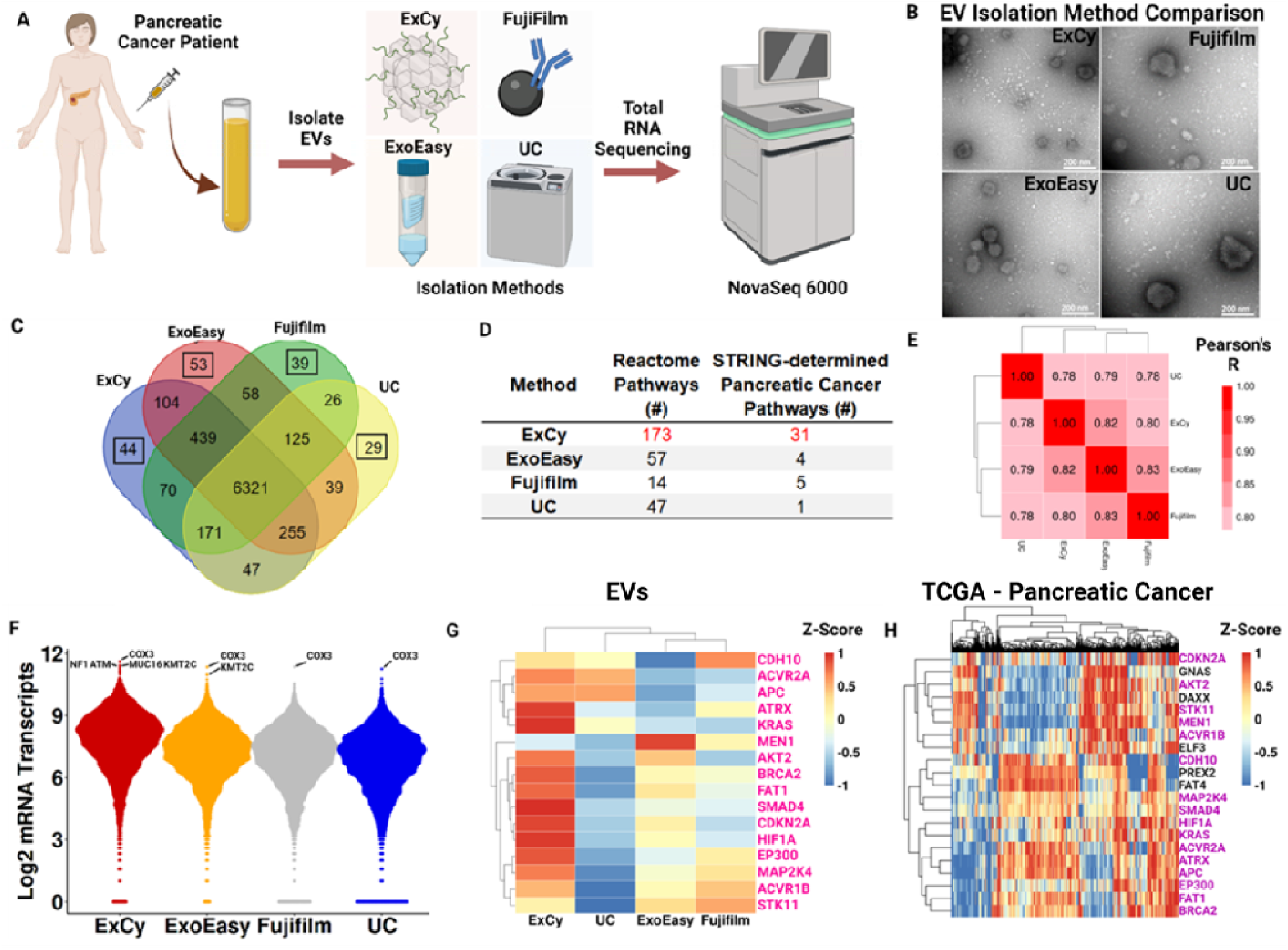
A patient zoomed-in exploratory analysis substantiates ExCy’s EV isolation quality. **A**. Illustrated scheme depicting zoomed-in patient analysis. 1 mL patient (Adenocarcinoma, pT1pN0, female, white) plasma was used. **B**. Transmission electron microscopy showing EV morphology across the four methods. **C**. Venn diagram illustrates mRNA comparison between isolation methods from total RNA sequencing. Highlighted box shows the number of unique mRNA annotations. **D**. Pathway analyses, using gProfiler and padj <0.05, on the unique mRNA annotations across each method. STRING-associated pancreatic cancer pathways were obtained from determined Reactome pathways. **E**. Method-by-method correlation matrix comparing similarity in mRNA profiles across methods. **F**. Beeswarm plots depicting mRNA transcript distributions; top mRNA transcripts were annotated, followed by filtering for pancreatic cancer associations using TCGA’s Cancer Gene Consensus analysis. **G**. COSMIC heatmap for all mRNA transcripts across each method. Pink colored labels indicate if it also appeared in the TCGA set. **H**. COSMIC heatmap using TCGA’s curated pancreatic cancer database to define patient baseline mRNA transcript level and profile to correlate with EV isolated transcript profiles. Purple colored labels indicate if it also appeared in the isolated EV set.

Next, we performed exploratory analysis to assess per-patient mRNA differences across the EV isolation methods (Figure 4, Figures s14-17, Supplementary File 1). Based on this analysis, we selected a representative tumor patient (Figure 4A) to show each method isolated representative EVs by TEM imaging (Figure 4B) and average size quantification (Figure s17), indicating a disparity in the EV population. Next, we visualized the similarities and differences in annotated mRNA across each method (Figure 4C), followed by pathway analysis with gProfiler on the unique mRNA annotations (Supplementary File 3) to probe whether the isolation-specific signatures indicate contribution to known pancreatic cancer pathways. To do so, we tabulated all the statistically significant Reactome ^88^ pathways from gProfiler (Supplementary File 4), then cross-referenced against STRING ^89^ to find Reactome-associated pancreatic cancer pathways (Figure 4D; Supplementary File 5). Given the number of pancreatic cancer pathways found, we further posited if the methods were enriching for pancreatic cancer related markers. Figure 4F shows the top mRNA transcripts across the methods, in which COX3, a prognostic marker in pancreatic cancer ^63, 90, 91^, appeared unilaterally. To find if the top mRNA transcripts could be cancer associated, we applied TCGA’s Cancer Gene Census analysis, which queries transcripts against the Catalogue of Somatic Mutations in Cancer (COSMIC)^92^, to find cancer-associated transcripts, NF1, ATM, MUC16, and KMT2C were found at higher mRNA levels with ExCy.

Since all isolation methods may pull down EVs with the similar cancer-related mRNA transcript levels, we also queried COSMIC ^92^ for pancreatic cancer related mRNA transcripts to visualize differences in their mRNA transcript levels between EV isolation methods (Figure 4G). Notably, ExCy identified more cancer related mRNA transcripts compared to the other methods (Figure 4G and Figure s16). We furthered compared the EV COSMIC mRNA transcript levels to TCGA pancreatic cancer data (Figure 4H) to understand what the relative difference between EV and TCGA transcripts observations were. ExCy showed obtaining much more COSMIC-associated mRNA transcripts than the other methods and was more representative of the TCGA profile.

### 3.4. Landscaping functional pathways analysis of isolated EVs

We posited that if method-dependent EV-mRNA transcripts lead to confounding or conflicted conclusions, could we understand the disparity level between methods for biomarker discovery in pancreatic cancer. To address this question, we performed pathway analysis with gProfiler on the unique mRNA annotations in each tumor patient across all methods relative to cancer and pancreatic cancer pathways (Supplementary file 5), then tabulated the unique and shared pathways per method across all tumor patients (Figure 5A). Each method contains distinct cancer-related pathways, but the shared pathways (i.e Translation, GPCR ligand binding, GPCR downstream signaling, and Disease) for the methods indicated different mRNA transcripts contributing to the pathway’s regulation. We further analyzed the pancreatic cancer association of the method-specific mRNA transcripts in the shared pathways by using TCGA’s Cancer Gene Consensus analysis on each pathway to determine pancreatic cancer significance.

**Figure 5.**
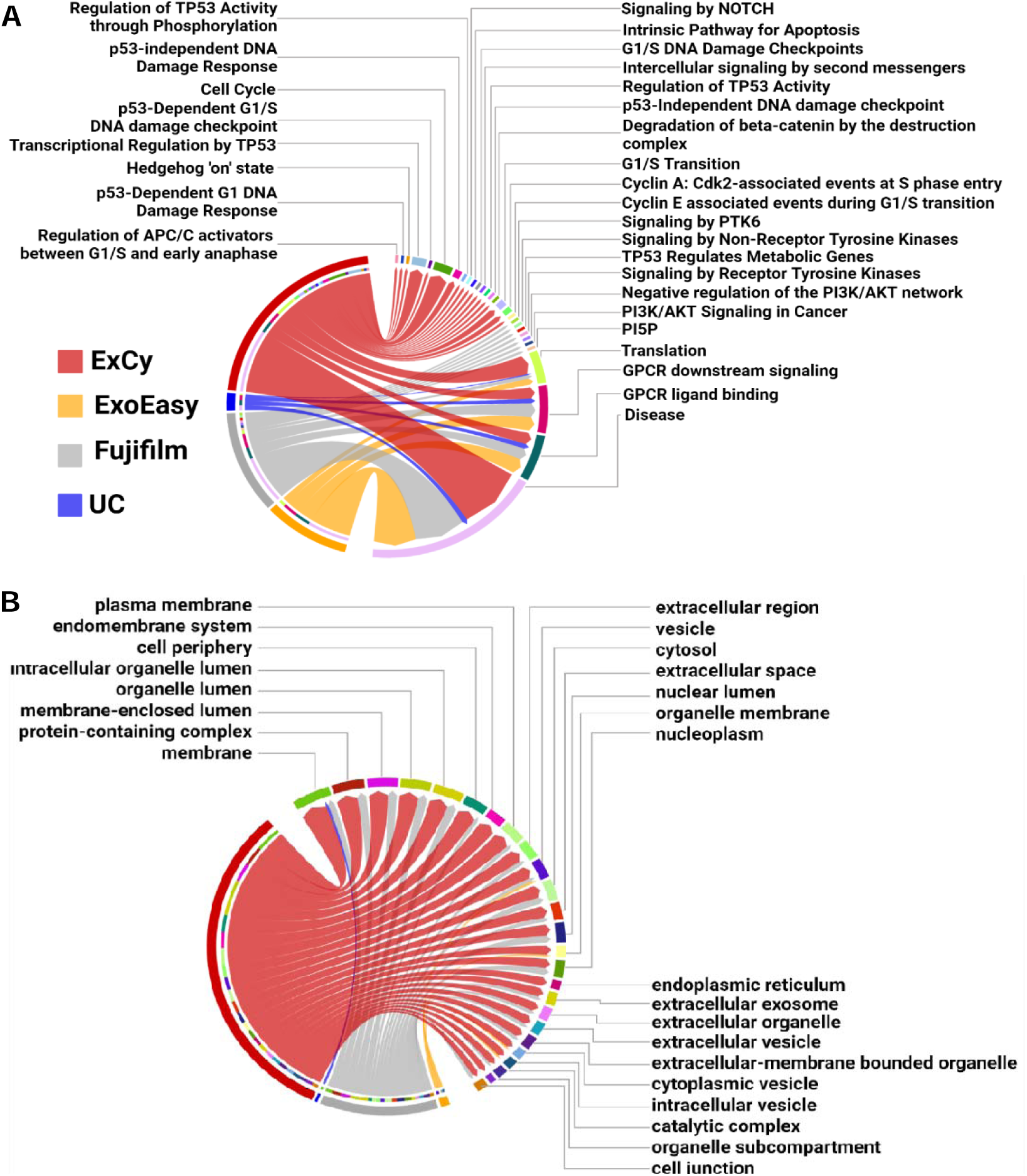
ExCy obtains statistically more EV-related and specific pathways that indicate the cancer and pancreatic cancer pathways from exploratory analysis. **(A)** Chord plot illustrating, at an exploratory level, pancreatic cancer related pathways associated to method dependent isolated EVs. The edges in the chord plot represent the number of annotated mRNAs found in each pathway. **(B)** An additional chord plot illustrating the top 25 statistically significant cellular component pathways found by gProfiler after separate differential analysis between EV isolation methods by edgeR on pancreatic cancer patients (N=4) and healthy control (N=3), then ranking differential transcripts to each isolation method across all patient samples (N=7).

The only overlapping pathways that were assessed by TCGA’s Cancer Gene Consensus analysis to have pancreatic cancer association from EV mRNAs was “Disease”, and only ExCy and Fujifilm’s mRNA transcripts in the Disease pathway were revealed by TCGA’s Cancer Gene Consensus analysis to be more likely mutated: MUC4 and CDK4 and CD79A and ARAF, respectively. To further validate if those mutated genes could come from non-EV contamination, we tabulated all differential transcripts in the healthy and tumor sample sets, then ranked the differential mRNA transcripts to an isolation method, and applied pathway analysis specifically for gene ontology, cellular component (GO:CC), because GO:CC is the pathway specific for extracellular vesicle ontology as shown in Figure 5B (Supplemental File 3). These results further provide support that ExCy is isolating transcripts related to many EV-specific terms, indicating high isolated EV homogeneity. In contrast, UC is only enriched for the membrane GO term (GO: 0016020). Since our differential analysis evaluates method-dependent mRNA transcript levels through statistical comparisons, and we further ranked the mRNA transcript relative to the isolation method, this result indicates that UC isolation has more technical bias. Given how extracellular vesicles have membrane profiles similar to that of parent cells, the EV transcriptomic profile could reveal their cellular origin. Thus, the results from ExCy isolated EVs showed the highest relevance to pancreatic tumors and originating cellular components, indicating the highest EV isolation quality, given the higher EV-related mRNA transcript levels when compared to ExoEasy, Fujifilm, and UC. ExCy beads are designed for high specificity with the NanoPom topography, enhancing the capture-release process with the insertion peptide’s selectivity for extracellular vesicles under acidification and removing non-EV particles by washing. The data supports that ExCy’s EV isolation is superior to the compared EV isolation methods, ExoEasy, Fujifilm, and UC, in terms of isolated EV purity.

### 3.5 Differential expression analysis between EV isolation methods

We analyzed differential expressions using edgeR (Figures 6A and 6D) to investigate the statistically significant differences in the method-dependent mRNA transcript levels between the healthy and pancreatic cancer patient plasma groups. In healthy patients, all methods appear to obtain statistically similar mRNA transcript levels, except for 103 mRNAs with differential expression from the hierarchical clustering analysis shown in Figure 6A. Notably, the differential expression pattern from the UC group is statistically different from ExCy, Fujifilm, and ExoEasy groups, while Fujifilm and ExoEasy did not statistically differ from each other (Figure s18). When compared to Fujifilm and ExoEasy, the ExCy group statistically differed for 3 mRNAs with significantly fewer transcript counts for PUF60 and VCX2 and more for SSX4. Therefore, using UC isolation as the standard benchmark may lead to false conclusions regarding an isolation method’s true capability to obtain high quality EVs. In consideration of each isolation method’s procedure, our observation also suggests the potential contamination by non-EV structures associated with UC and Fujifilm’s isolation. This observation is also consistent with our EQI determination in Figure 3F & G. Based on our findings, we ranked each method’s differential mRNA transcript levels within the healthy sample set (Figure 6B) and then plotted the top 10 ExCy-ranked mRNAs (Figure 6C) to understand further trending differences in mRNA profiles between the EV isolation methods. When we assessed ExCy’s top-ranked differential mRNAs against all other methods, we observed that ExCy was, on average, the most consistent to obtain mRNA transcripts (Figure 6C), in contrast to the UC or Fujifilm isolation method. Most importantly, we observed that ExCy consistently identified a significant EV marker, CAPN1. When queried by UniProtKB, CAPN1 corresponds to the Extracellular Exosome pathway^93^. It is worth mentioning that differences in EV isolation approach are a strong signal in the resultant data that may confound biological conclusions.

**Figure 6.**
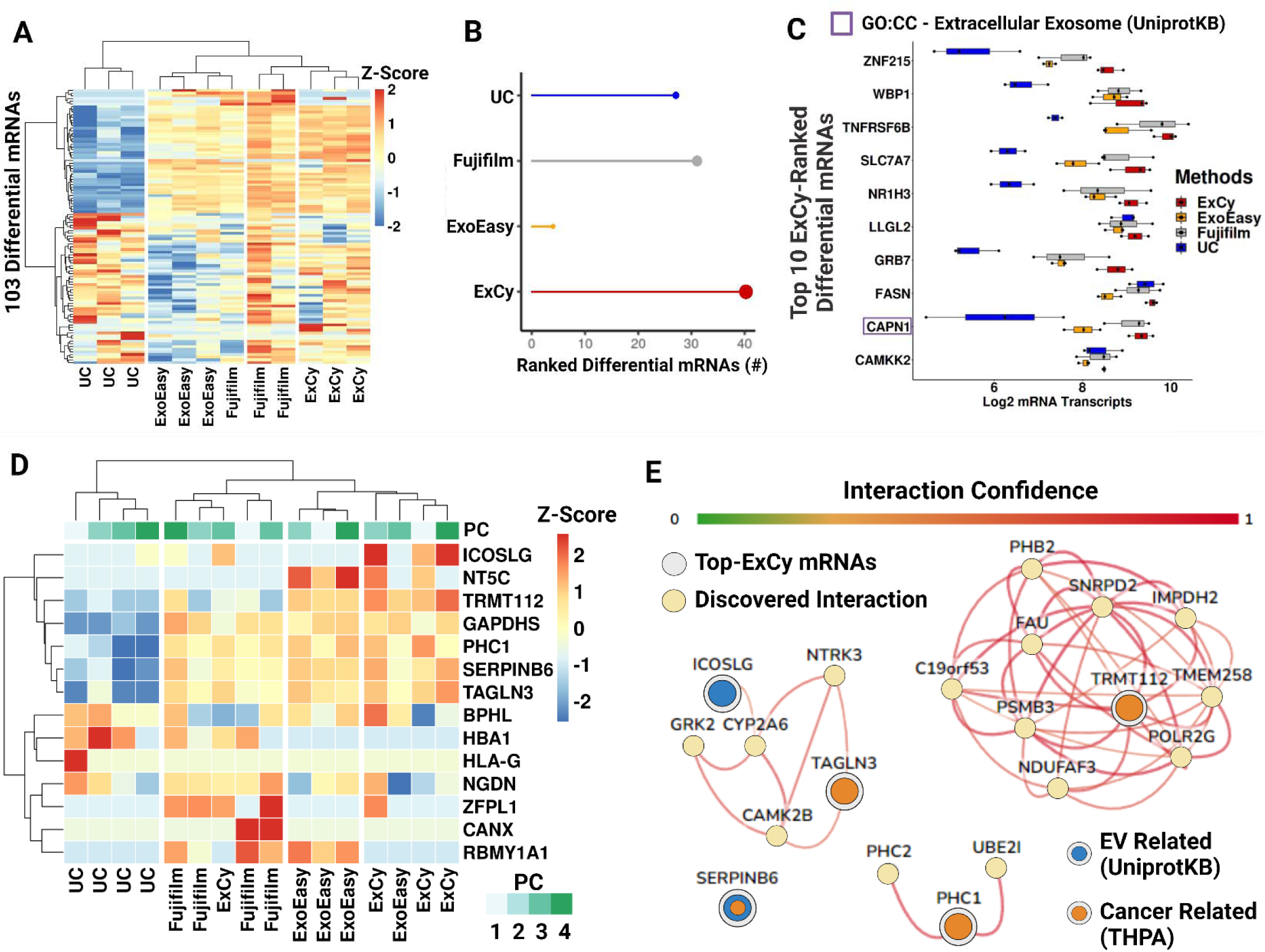
Differential expression analysis of mRNA profiles derived from different EV isolation methods. **A** Heatmap showing the landscape of differentially expressed transcripts in healthy patient plasma isolated EVs across different isolation methods, as determined by edgeR (FDR < 0.05). The color gradient indicates the Z-score. Hierarchical clustering was performed using Ward’s D2 method^94^. **B**. Differential transcripts in healthy patients were ranked across each method based on their average transcript count. The filled point represents the total number of differentially ranked mRNAs in the isolation method, while the line segment helps visualize the relative difference of total differentially ranked mRNAs between methods. **C**. ExCy’s top 10 ranked mRNA transcripts compared to other EV isolation methods illustrated by the box-and-whisker plot. **D**. Heatmap showing the landscape of differentially expressed genes in pancreatic cancer patient plasma isolated EVs across different isolation methods, as determined by edgeR (FDR < 0.05). **E**. ExCy ranked mRNA transcripts were inputted into HumanBase’s GIANT global tissue network. Input mRNA transcripts were further identified as EV related by UniprotKB or cancer related by The Human Protein Atlas, while the connections between nodes represent the interaction confidence greater than 0.60.

Next, we investigated the tumor cohort (Figure 6D) and observed a smaller number of differentially expressed genes between each EV isolation method. As observed consistently with the healthy sample cohort, UC derived transcriptomic profile significantly differs from ExCy, ExoEasy, and Fujifilm groups which showed similarity in between. Next, we extracted all ExCy ranked differential tumor transcripts and performed functional enrichment analysis using the HumanBase enrichment tool and global functional interaction network (Figure 6E). Result identifies ICOSLG gene from ExCy isolated EVs to show interaction relationship with CYP2A6 which is a critical network of drug metabolism for monitoring drug responses from patients undergoing treatment (Figure 6E). However, this interaction may not be detected using ExoEasy, Fujifilm, or UC isolated EVs to quantify mRNA transcript levels. Interestingly, TAGLN3, a potential oncogene and regulator of RNA Polymerase II, was detected by ExCy, ExoEasy and Fujifilm, but not by UC. This observation suggests that a potential tumor biomarker must be associated with EV-biology in a given isolation since ExCy also isolated ICOSLG at high levels. Similarly, for TRMT112, neither Fujifilm nor UC may detect this transcript, missing TMRT112’s impact as an oncogene and subsequent role as a macromolecular methylator and rRNA effector. Finally, PHC1 and SERPINB6 were not detected by UC; however, their roles as either a pancreatic-cancer specific oncogene, regulating GMNN expression in the pancreas, or potential EV-associated oncogene, as a serine protease, respectively, further indicates that ExCy, ExoEasy, and Fujifilm could isolate EVs more consistently to detect tumor associated markers. In light of this observation, we further looked at the GMNN mRNA levels across the methods (Figure s19), indicating that ExCy, ExoEasy, and Fujifilm extract GMNN mRNA reliably, which would be expected based on observing PHC1 transcripts. However, ExCy had the highest average and least variable mRNA transcript level, suggesting further evidence for higher EV homogeneity. Our findings highlight the urgency of using an additional optimal EV isolation method to validate identified markers for functional analysis.

### 3.6 Differential expression analysis for exploring circulating markers associated with pancreatic tumor

We hypothesized that markers identified by the method must be associated with EV biology within the context of pancreatic cancer, which is testable through functional enrichment after differential analysis. Given the limited statistical power to discover tumor associated markers in each method^95–98^, we slightly adjusted the FDR from 0.05 to 0.10 to enable a wider pool of markers to be analyzed. We expected to observe considerable variation in identified method-specific markers. Notably, none of the differential markers in each method overlapped with the other methods (Figure 7A). Under the assumption that all methods obtain EV transcripts, we pooled all the markers together for pancreatic enrichment analysis by HumanBase to ascertain functional potential. At an interaction confidence of 0.60, HumanBase revealed that ExoEasy and UC’s identified markers did not interact with the pancreatic network. However, both ExCy and Fujifilm were revealed to have a marker showing strong interconnection in pancreatic tissue. Interestingly, a subnetwork within HumanBase remained present above the cutoff. When the interaction confidence was adjusted from 0.60 to 0.53, the subnetwork integrated into ExCy’s network, suggesting ATP6V0B’s regulatory impact, which is much larger than the Fujifilm-identified biomarker BAG6.

**Figure 7.**
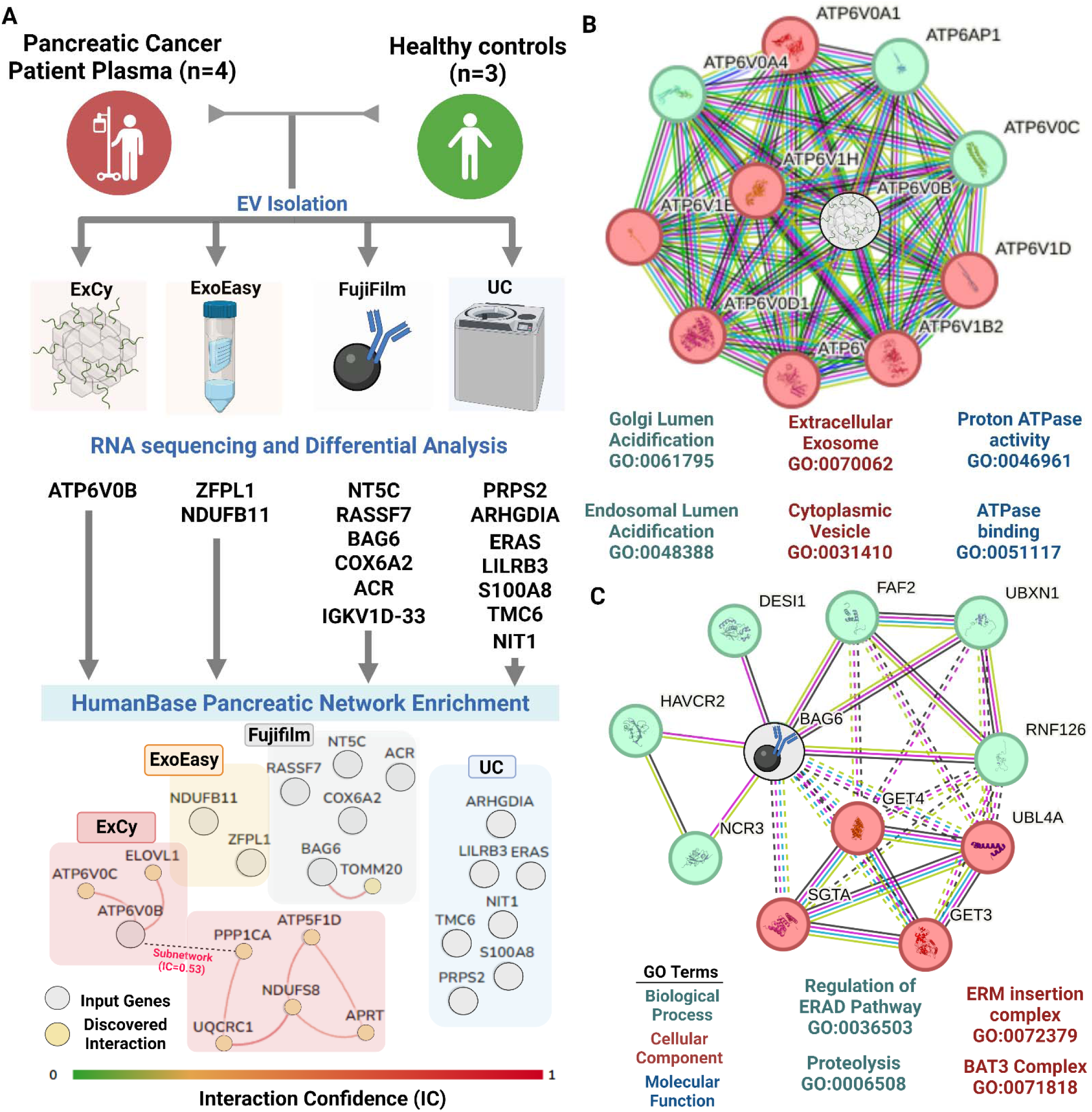
Differential expression analysis indicates ExCy’s capability to find relevant pancreatic tumor associated markers. **A** Differential analysis was applied to compare tumor versus healthy samples to find distinct markers (edgeR FDR <0.1) per method. We then pooled all the differential markers into the HumanBase GIANT pancreatic network for enrichment analysis to discover interactions within the pancreas. The connections between nodes represent the interaction confidence greater than 0.60. Only ExCy (ATP6V0B) and Fujifilm (BAG6) differential transcripts were deemed valid based on their connection within the network. **B**. We queried STRING on ATP6V0B from ExCy EV isolation to investigate protein interactions and their representative ontological terms as a tumor associated marker, and **C**. BAG6 from Fujifilm isolation method.

We further investigated those marker potential by looking at STRING’s functional protein network of ATP6V0B (Figure 7B) and BAG6 (Figure 7C) respectively. In STRING, we used an interaction confidence of 0.90 to ensure maximum confidence regarding the biological capacity. We extracted their gene ontology (Supplementary File 6), revealing that ExCy’s discovered marker, ATP6V0B, is a highly relevant EV marker that increases acidification (logFC = 2.02) of the endosomal and Golgi lumen in pancreatic tumors, dysregulating both EV production and cytosolic ion-channel gradients. This observation strongly supported that markers identified by an EV isolation method are associated with EV biology within the context of pancreatic cancer. On the other hand, Fujifilm’s discovered marker, BAG6, is a cytosolic molecular chaperone participating in delivering proteins to the endoplasmic reticulum or proteasome, unspecific to EV biology. In this view, ExCy was the only EV isolation method that successfully identified an EV-relevant marker that may translate into pancreatic cancer biology.

## 4. DISCUSSION

EVs are increasingly used to understanding regulatory mechanisms and their use in clinical research has resulted in clinical trial establishment ^6, 99^. However, each component of the EV pipeline contributes to the observed EV heterogeneity^6, 9, 22, 25, 26, 28, 30, 100–108^ and this heterogeneity has far-reaching implications, most notably in terms of the interpretations drawn from EV samples concerning biological processes and their reproducibility^9, 11, 25, 28, 34–40^. EV isolation induced heterogeneity can lead to conflicting conclusions and hinder the ability to replicate findings, thereby impeding the overall progress of EV research. We first addressed EV heterogeneity at the isolation level by developing a novel strategy to isolate EVs using an insertion peptide enabled capture-release process, which can enhance the selectivity for EV isolation. Due to the well-observed selectivity to insert into unilamellar particles^51–56, 81^, in addition to ExCy’s unique NanoPom topography^41^. The insertion peptide enabled capture-release EV isolation can reduce contamination by lipoproteins and nano-debris. Considering the insertion peptide energetics within liposomal systems, smaller size nanoparticles will have higher surface curvatures, thereby conferring substantially larger surface energy^109^. Thus, the insertion peptide’s association to smaller EV surfaces and subsequent capture could be thermodynamically favorable compared to their larger sized counterparts such as cells.

We next created a statistical algorithm standard for comparison between EV isolation methods by employing data science techniques to define the extent of EV heterogeneity, which we demonstrated in human plasma samples. Using this framework, we established a statistical benchmark for the expected quality of EV isolates, demonstrating how standardization was essential for direct comparison between isolation methods. Such data also supports that our developed ExCy isolation approach enables broad applications for isolating EVs from various biological fluids, which contributes to efficient EV isolation for reliable EV research. Moreover, an additional optimal EV isolation method is crucial in biomarker discovery to validate identified markers for functional analysis.

Direct comparison between isolation methods are essential in the field. We developed a statistical metric for comparing 4 different total EV isolation methods (ExCy, ExoEasy, Fujifilm, and UC). Though the EV isolation is for total EV selection, the isolation mechanism differs entirely between isolation methods. ExCy captures EVs by transmembrane insertion, ExoEasy isolates by membrane affinity and centrifugation, Fujifilm by phosphatidylserine affinity, and UC by density, resulting in distinct EV isolate profiles. MISEV provides existing alternate semi-quantitative metrics to investigate EV quality, including particle-to-protein and particle-to-RNA; however, even when considering the same source, these metrics are confounded by the isolation method. Therefore, we adjusted for method-specific technical bias using quantile-quantile normalization to adjust their resulting measurements. Quantile transformation rescales measurements between 0 and 1 based on the cumulative distribution function, defining all values equally likely. Allowing an equal likelihood of measurement, when given the same EV source, enables comparison of different EV isolation methods (Figure 3). When we applied the quantile transformation, we did not observe any loss of information. Instead, we demonstrated that our approach coincided with the field’s understanding that EV isolates are still method dependent. We also demonstrated that comprehensive characterization is required to indicate an EV isolate’s quality before subsequent translational analysis and that relying on paired measurements has likely obfuscated potential discoveries for confounding existing conclusions reported in the field. To address this issue, we laid a foundation for characterizing an EV isolate’s quality by demonstrating that the EQI could reveal a method’s expected EV isolate homogeneity upon resampling. If we included the studies from EV-TRACK, our statistical metric may enable EV quality comparisons across EV isolation methods and substantially improve the overall rigor in the field of EV research.

We validate ExCy’s EV isolation capability through next generation sequencing. We confirmed this premise by showing the differences in RNA transcript quality through the FREYA pipeline with HISAT2 (Supplementary Figure 8) and FastQC (Supplementary File 2). EV transcriptomics are subject to confoundment regarding both EV heterogeneity and transcriptomic practices. Using FastQC, we showed the significant difference in library quality after processing. We observed differences in the total RNA and mRNA distribution between the EV isolation methods; however, the correlated mRNA transcript levels between all methods were moderate (Pearson’s r = 0.78, Supplementary Figure 13). On the other hand, we showed through differential analysis that all methods are statistically similar in the bulk of their EV isolate mRNA transcript levels, differing by a distinct fraction of mRNA transcripts, though this observation may be affected by the limited sample power in our study. Based on the mRNA transcript levels, pathway analysis, and differential analysis, we showed that ExCy may be the most useful to developing applications. Additionally, from an EV quality standpoint, we also show that the EQI indicates that ExCy, relative to ExoEasy, Fujifilm, and UC, obtains more homogenous EV isolates upon resampling (EQI = 79.78% vs 69.76% geometric average of other methods, Figure 3); However, given the limited sample power and number of assays conducted to measure total EV yield, RNA, protein, size and size distribution in the dataset, we suggest that scaling both the sample size and total assay number may fully reveal the expected EV isolate quality from an isolation method provides.

We also proposed that EV-isolation method-specific markers in disease contexts must need to be associated with EV biology, which we demonstrated through functional enrichment after differential analysis. Total EV isolation methods are expected to pull down similar EV transcripts, given that these methods are EV selective; however, we showed that regardless of isolation technique, the expected EV quality strongly affects differential analysis to find relevant markers (Figure 7A), since no overlap in differential markers were identified. Although there is limited sample power to find tumor associated markers, based on the EQI (Figure 3), it may be unlikely that UC and Fujifilm would be consistent in their EV resampling capability, which leaves significant uncertainty regarding their identified markers. To tackle this uncertainty, we cross-validated HumanBase’s enrichment analysis with STRING’s functional protein networks, showing that ExCy’s identified marker was highly significant for EV biology in the pancreatic cancer disease context, which proves our hypothesis. In the next generation sequencing analysis, we suggest increasing the technical and biological replicates, and include an EV isolation control to support translational claims.

In summary, we developed a method to isolate EVs combining our established NanoPom topography with peptide-based capture and release, called ExCy. We also addressed the challenge regarding direct comparison between EV isolation methods by developing a statistical metric (ExoQuality Index, EQI) to directly quantify EV heterogeneity, which we validated at a transcriptomic level. Importantly, while we demonstrate all methods isolate EVs in some capacity, EV isolation methods should be controlled with another isolation technique to avoid conflicting outcomes, and UC is unable to serve as the reference or benchmark method, revealing itself as an outlier in this study. In this outlook, ExCy’s utilization in broad EV research as an isolation method or control could provide the least confounding evidence to support biological conclusions.

## Data and software availability

All raw and processed sequencing data generated by this study is available at the NCBI Gene Expression Omnibus (GEO; https://www.ncbi.nlm.nih.gov/geo/query/acc.cgi?acc=GSE246925). Computer code used in this manuscript is available at github.com/zfg2013/ExCy. All code used in this manuscript is shared at this repository, and to ensure reproducibility, we provide a README with version information for each tool plus any parameter settings used in the data processing and analysis.

## Acknowledgements

The authors would like to thank the Interdisciplinary Center for Biotechnology Research (ICBR) core facility at the University of Florida for sequencing services. This project is supported by NIH NIGMS MIRA Award 1R35GM113794, CFF HE21I0, and UFHCC GU pilot to Dr Mei He, and University of Florida Health Cancer Center Pilot Grant # AI-2022-02 awarded to Dr. Kiley Graim and Dr. Mei He. Corresponding contacts: &

## Supporting Information

Supplementary Materials – Supplementary Figs. 1-19

Supplementary File 1 – ExoQuality Index Dataset and Sequencing information. The dataset is used to compute the EQI and sequencing information regarding HISAT2’s RNA annotations per sample across all methods before and after Vesiclepedia mapping.

Supplementary File 2 – FastQC reports – External quality reports using FastQC, before and after applying trimmomatic, detailing each EV isolation’s transcriptomic sample assembly right after sequencing by the NovaSeq 6000. Each report will present basic assembly statistics followed by listing quality metrics including, Per base sequence quality, Per tile sequence quality, Per sequence quality, Per base sequence content, Per sequence GC content, Per base N content, Sequence Length Distribution, Sequence Duplication levels, Overrepresented sequences, and adaptor content.

Supplementary File 3 – gProfiler’s enrichment analysis. Enrichment analysis was applied to each EV isolation’s transcriptomic sample based on their unique mRNA annotations.

Supplementary File 4 – EV Isolation method-specific mRNA annotations and subsequent Reactome pathway. Supplementary File 5 – Patient STRING pathways and intersecting mRNA annotations.

Supplementary File 6 – STRING pathway analysis on BAG6 and ATPV06B

