## Supplemental File for "Peptide-based capture-and-release purification of extracellular vesicles and statistical algorithm enabled quality assessment"

* corresponding contact

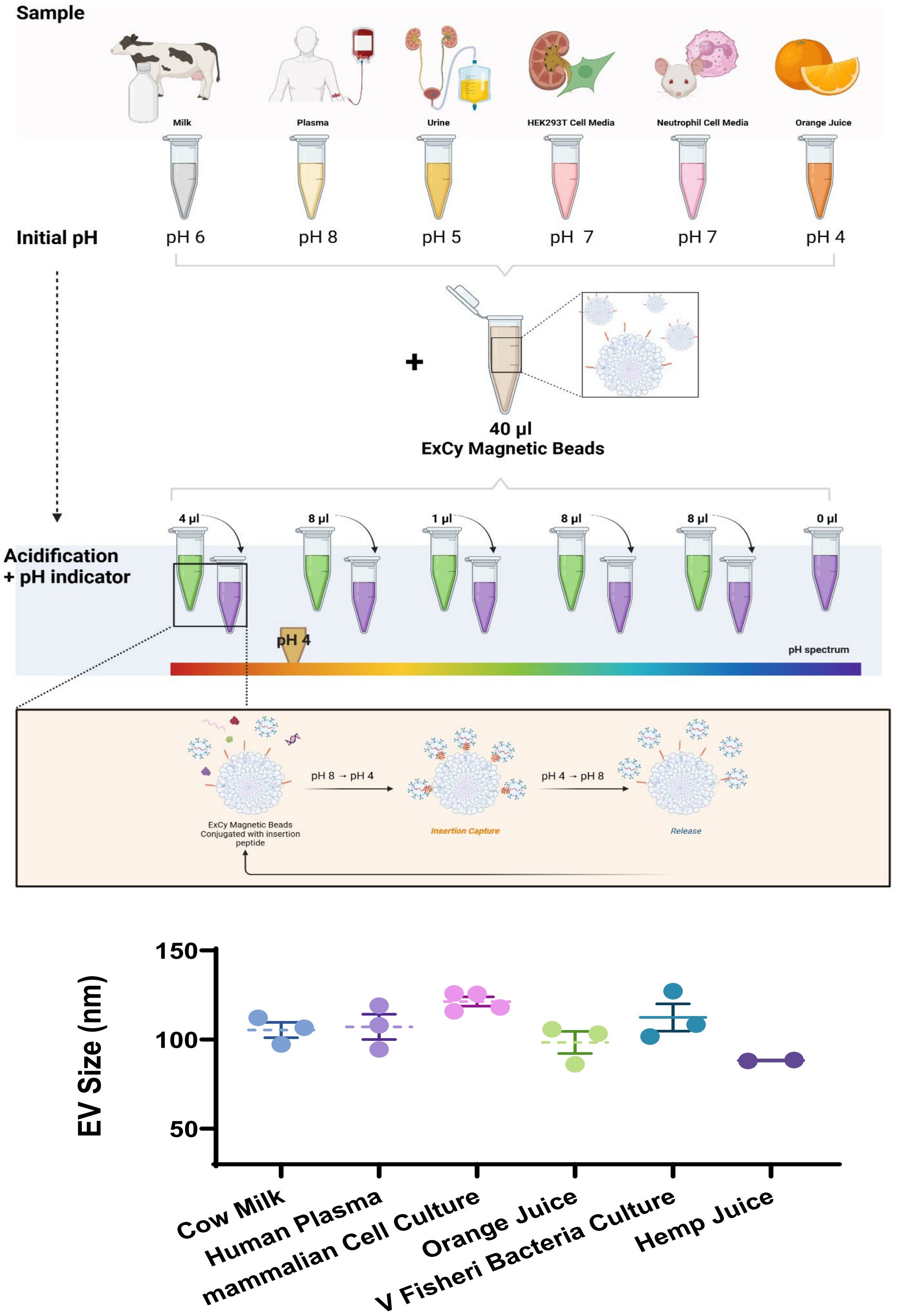


**Figure s1.** Broad workflow applicability of ExCy in various biological media with characterized EV size.

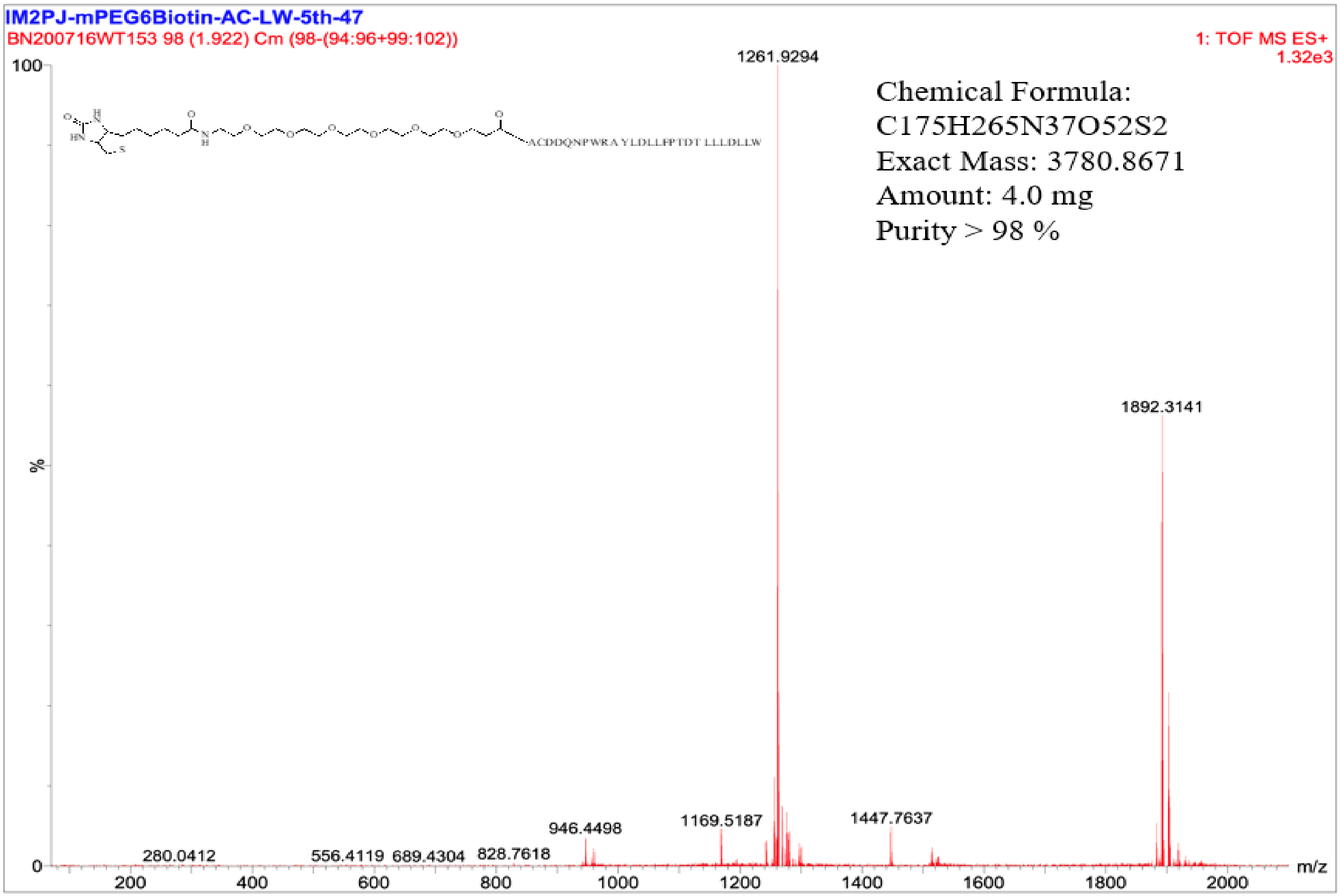


**Figure s2.** The mass spectrometric characterization and QC after peptide microwave synthesis.

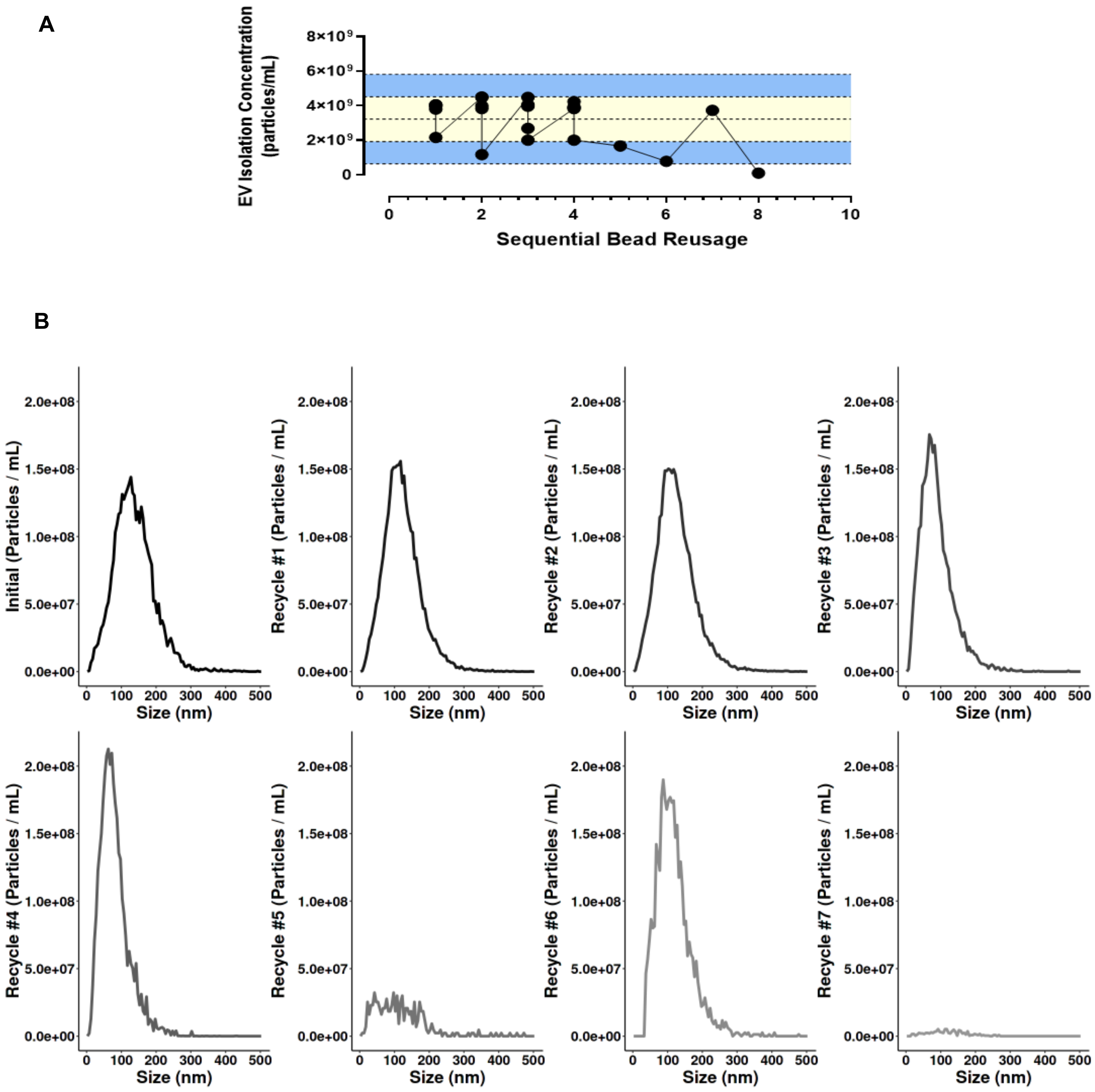


**Figure s3.** ExCy's capability to recapture EVs. **A)** EV isolate concentrations after ExCy's sequential usage in the same sample type (plasma), indicating limits of isolation reproducibility. Yellow bars indicate confidence interval, while blue bars fall out of range. **B)** Nanoparticle tracking analysis for each sequential isolation. Initial designates the starting EV isolate concentration from human plasma.

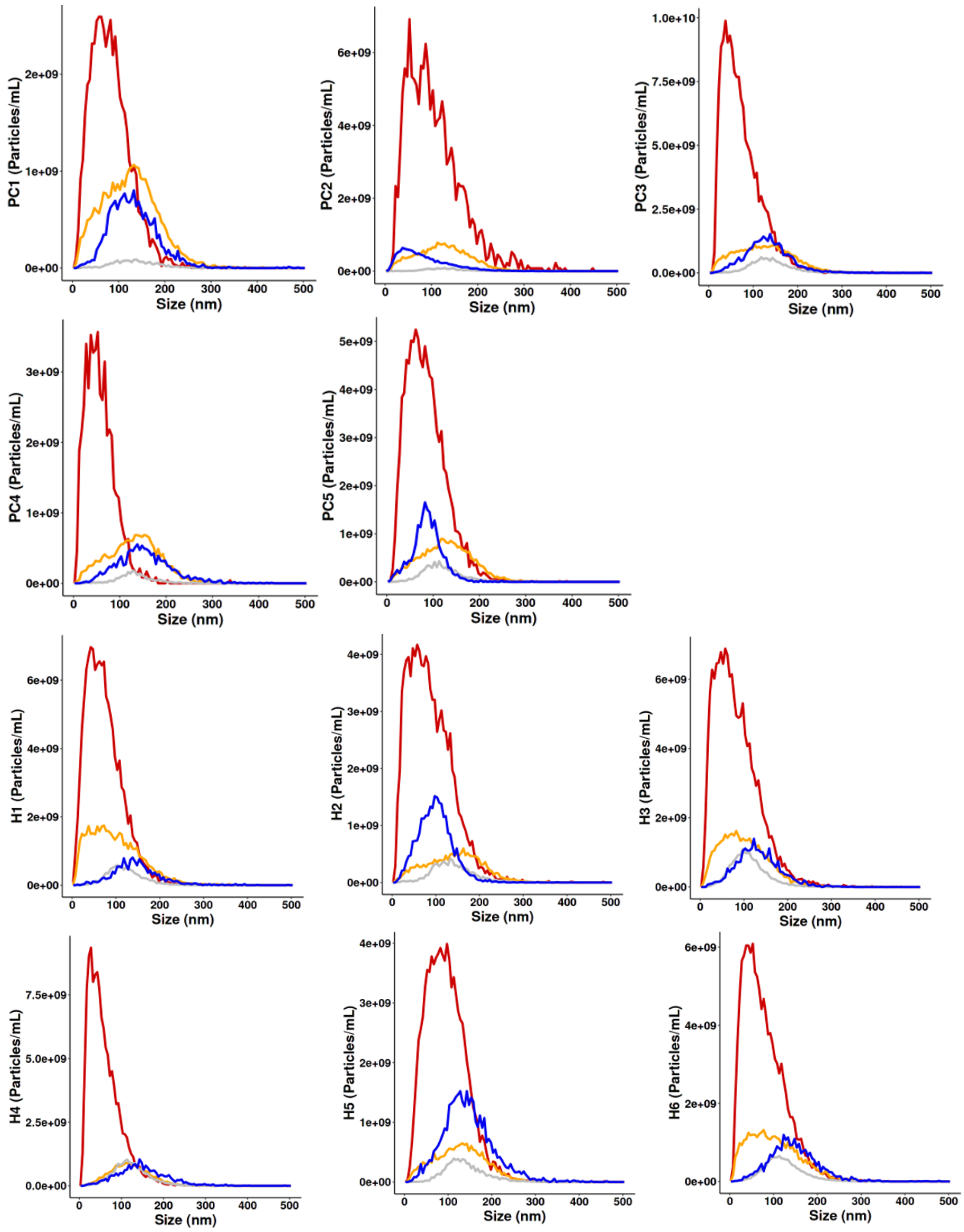


**Figure s4.** NTA size distribution profile from all human plasma samples from **A)** Pancreatic cancer and **B)** Healthy controls. Red line: ExCy; Blue line: UC; Orange: ExoEasy; Gray: FujiFilm.

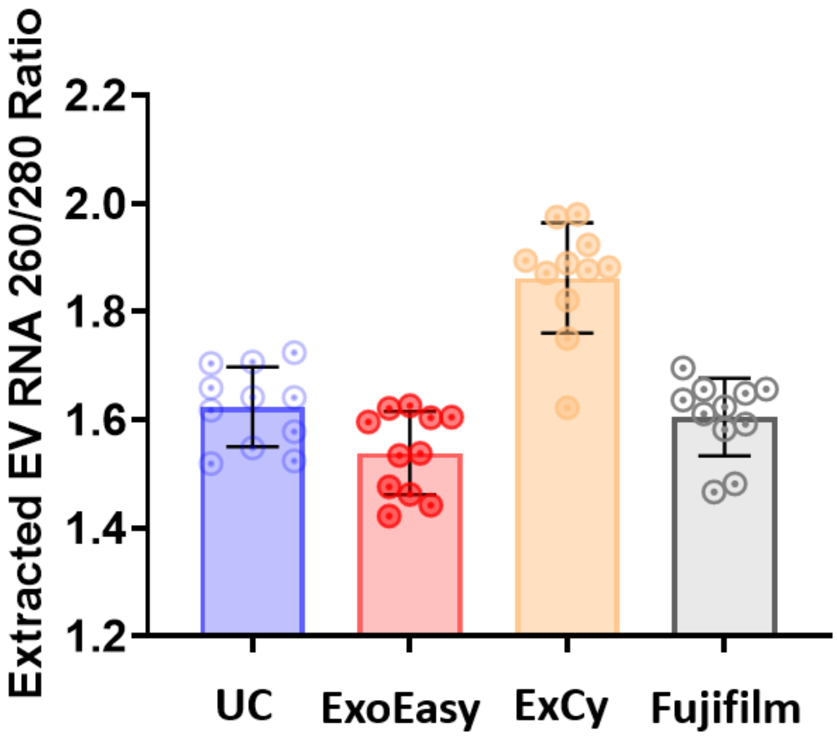


**Figure s5**. 260/280 nm$^{-1}$ analysis to examine RNA purity after applying Qiagen’s miRNeasy kit to extract total RNA.

**A)**

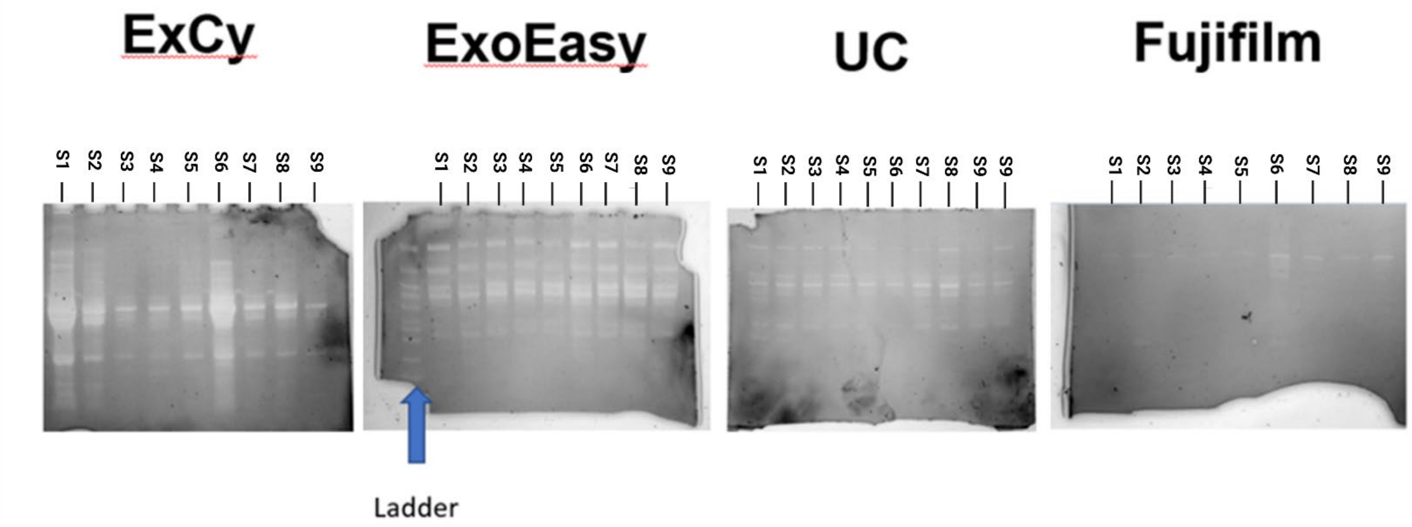


B)

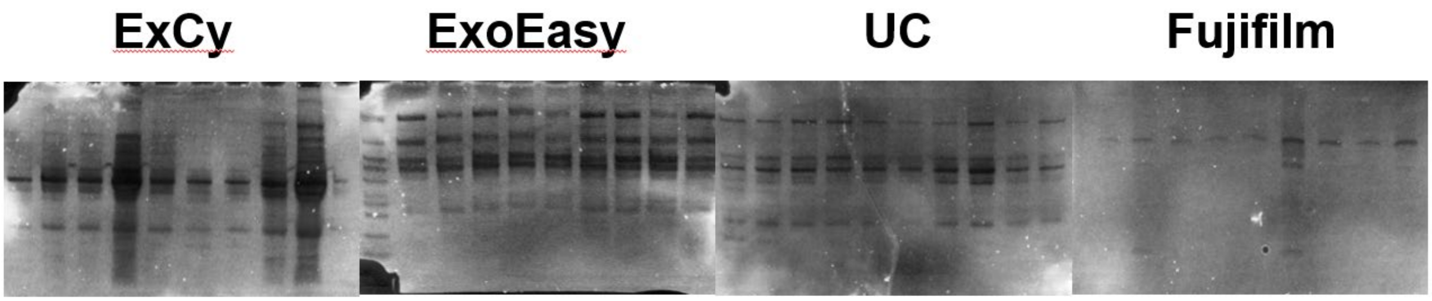


**Figure s6.** Protein gel analysis by Coomassie blue staining through BioRad imaging, within a tetra system, on the same samples for the four EV isolation methods. **A)** Unprocessed raw gel. S9 was added twice to UC, out of random selection, to assess protein gel variation. **B)** ImageJ processed**.**

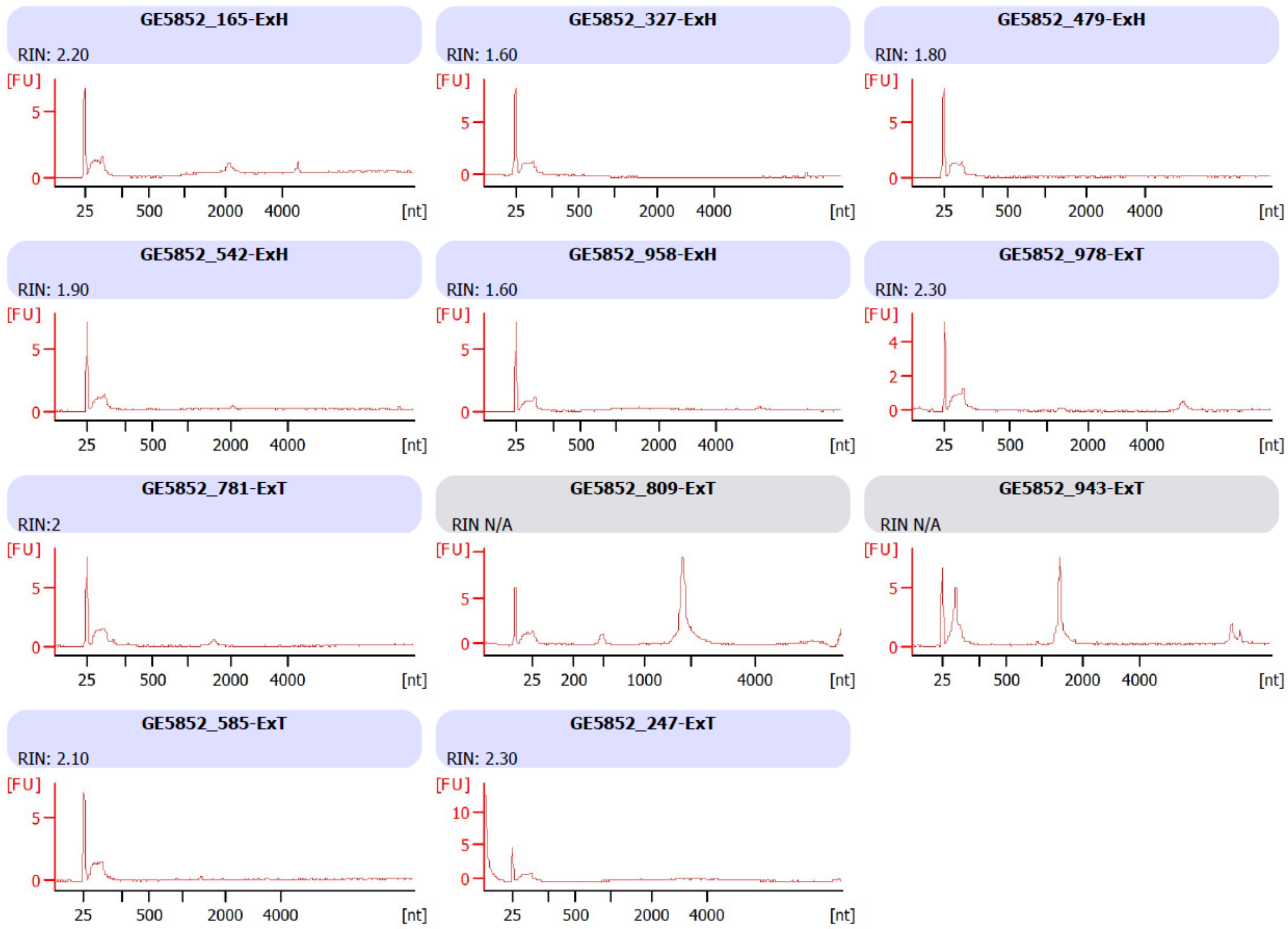


**Figure s7.** Bioanalyzer analysis on UC extracted RNA

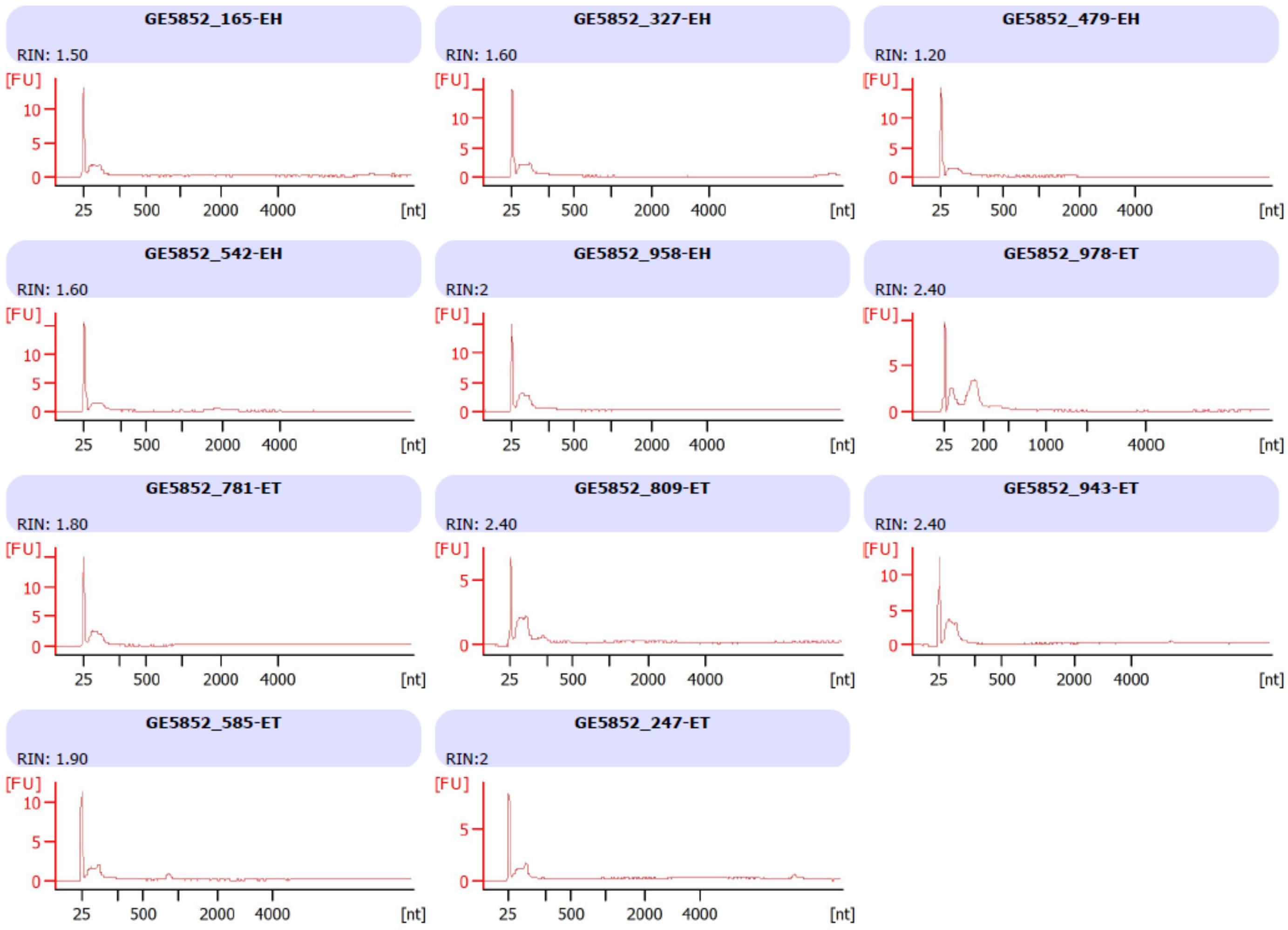


**Figure s8.** Bioanalyzer results on ExoEasy extracted RNA

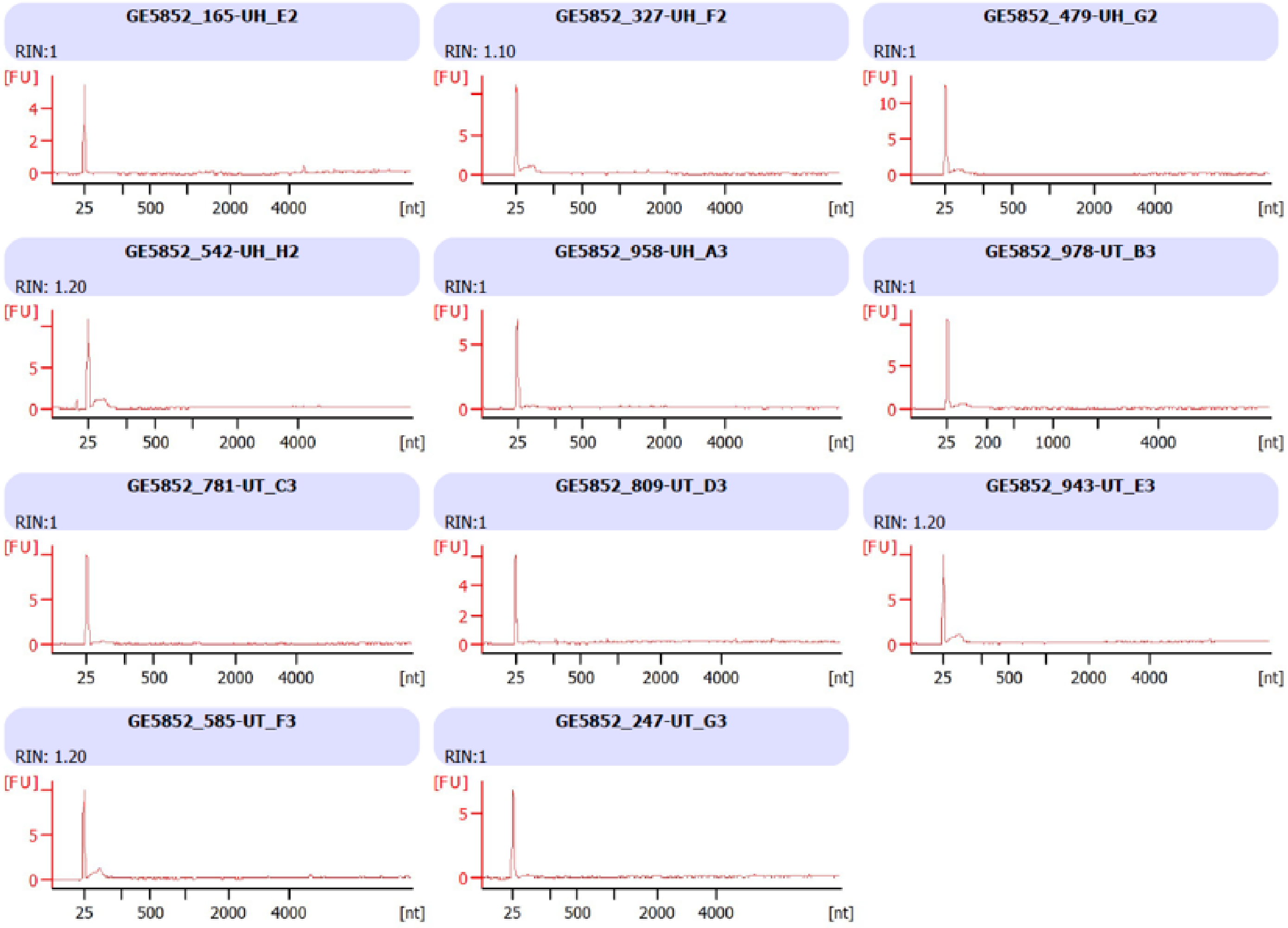


**Figure s9.** Bioanalyzer results on ExCy extracted RNAs

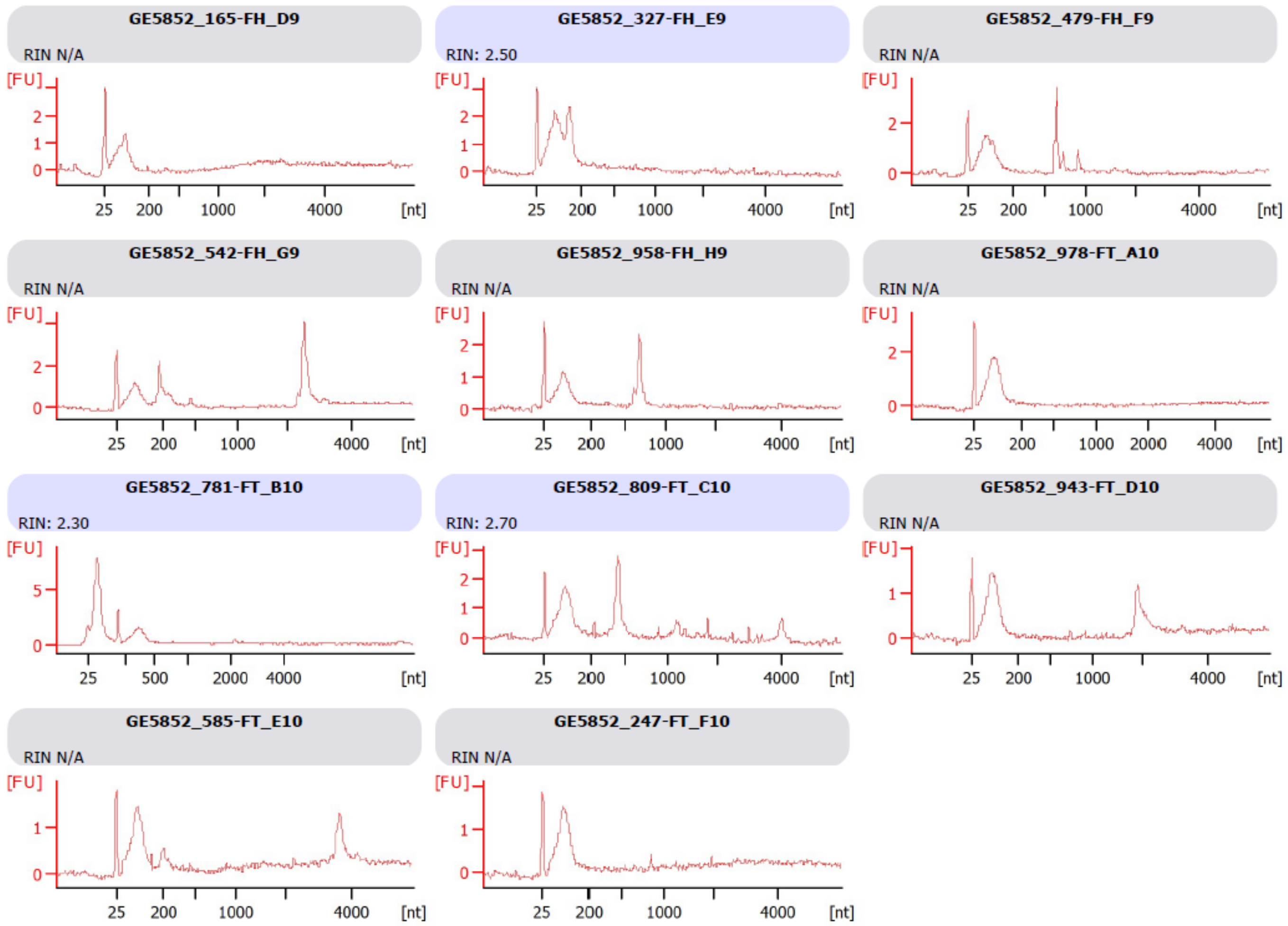


**Figure s10**. Bioanalyzer results on Fujifilm extracted RNAs

| Patients | ExCy | ExoEasy | FujiFilm | UC |
|---|---|---|---|---|
| **PC1** | 75.98 | 71.06 | 68.91 | 61.67 |
| **H1** | 80.94 | 78.84 | 89.93 | 60.74 |
| **H2** | 86.38 | 81.31 | 77.37 | 71.15 |
| **PC2** | 79.21 | 57.3 | 60.04 | 57.89 |
| **PC3** | 73.36 | 57.38 | 67.28 | 61.93 |
| **H3** | 77.34 | 78.23 | 74.23 | 62.46 |
| **PC4** | 68.09 | 73.53 | 79.91 | 67.74 |
| **AVG ± STDEV:** | 77.33 ± 5.79 | 71.09 ±10.00 | 73.95 ± 9.71 | 63.37 ±4.50 |

**Figure s11**. HISAT2 mapping rate across each method for all the patients.

### ExCy

| Patients | Count | AS | LINC | Intronic | lncRNA | snRNA | tRNA | Divergent | miRNA | NE-MT | OT | rRNA | R60-Y | Small NF90 | snoRNA | Variant U1 |
|---|---|---|---|---|---|---|---|---|---|---|---|---|---|---|---|---|
| PC1 | 21788 | 683 | 1137 | 31 | 174 | 8 | 0 | 136 | 375 | 0 | 7 | 0 | 1 | 0 | 86 | 3 |
| H1 | 23725 | 753 | 1217 | 40 | 203 | 6 | 0 | 163 | 419 | 0 | 7 | 0 | 1 | 0 | 83 | 3 |
| H2 | 24107 | 1216 | 46 | 202 | 10 | 0 | 157 | 475 | 0 | 7 | 0 | 1 | 0 | 96 | 6 | 0 |
| PC2 | 24016 | 771 | 1225 | 39 | 202 | 12 | 0 | 157 | 452 | 1 | 7 | 0 | 2 | 0 | 82 | 8 |
| PC3 | 21954 | 685 | 1112 | 36 | 187 | 6 | 0 | 141 | 335 | 0 | 7 | 0 | 0 | 0 | 68 | 2 |
| H3 | 22335 | 681 | 1157 | 35 | 187 | 3 | 0 | 148 | 391 | 0 | 7 | 0 | 0 | 0 | 89 | 2 |
| PC4 | 21384 | 666 | 1091 | 32 | 194 | 8 | 0 | 141 | 360 | 0 | 6 | 0 | 1 | 0 | 63 | 5 |

### ExoEasy

| Patients | Count | AS | LINC | Intronic | lncRNA | snRNA | tRNA | Divergent | miRNA | NE-MT | OT | rRNA | R60-Y | Small NF90 | snoRNA | Variant U1 |
|---|---|---|---|---|---|---|---|---|---|---|---|---|---|---|---|---|
| PC1 | 21108 | 653 | 1049 | 24 | 187 | 6 | 0 | 135 | 276 | 0 | 7 | 0 | 0 | 0 | 62 | 4 |
| H1 | 22262 | 686 | 1140 | 31 | 189 | 6 | 0 | 151 | 290 | 0 | 7 | 0 | 0 | 0 | 74 | 4 |
| H2 | 19490 | 571 | 968 | 25 | 160 | 1 | 0 | 117 | 223 | 0 | 5 | 0 | 0 | 0 | 56 | 1 |
| PC2 | 19349 | 584 | 954 | 20 | 166 | 5 | 0 | 120 | 289 | 0 | 6 | 0 | 1 | 0 | 65 | 0 |
| PC3 | 18503 | 548 | 916 | 21 | 155 | 5 | 0 | 120 | 258 | 0 | 6 | 0 | 0 | 0 | 45 | 4 |
| H3 | 20864 | 640 | 1072 | 28 | 164 | 4 | 0 | 132 | 288 | 0 | 7 | 0 | 0 | 0 | 62 | 3 |
| PC4 | 22509 | 701 | 1149 | 33 | 177 | 5 | 0 | 145 | 380 | 0 | 7 | 0 | 1 | 0 | 92 | 4 |

### Fujifilm

| Patients | Count | AS | LINC | Intronic | lncRNA | snRNA | tRNA | Divergent | miRNA | NE-MT | OT | rRNA | R60-Y | Small NF90 | snoRNA | Variant U1 |
|---|---|---|---|---|---|---|---|---|---|---|---|---|---|---|---|---|
| PC1 | 20129 | 633 | 1010 | 21 | 176 | 6 | 0 | 130 | 304 | 0 | 5 | 0 | 0 | 0 | 63 | 1 |
| H1 | 25408 | 806 | 1265 | 45 | 210 | 14 | 0 | 175 | 519 | 0 | 7 | 0 | 1 | 0 | 94 | 8 |
| H2 | 20692 | 600 | 1042 | 32 | 177 | 7 | 0 | 133 | 332 | 0 | 5 | 0 | 0 | 0 | 60 | 3 |
| PC2 | 19663 | 591 | 991 | 25 | 169 | 2 | 0 | 133 | 266 | 0 | 7 | 0 | 0 | 0 | 57 | 0 |
| PC3 | 21528 | 676 | 1095 | 30 | 185 | 8 | 0 | 138 | 327 | 0 | 7 | 0 | 2 | 0 | 73 | 3 |
| H3 | 20592 | 610 | 1056 | 27 | 178 | 4 | 0 | 135 | 251 | 0 | 6 | 0 | 0 | 0 | 65 | 2 |
| PC4 | 23009 | 737 | 1167 | 34 | 195 | 4 | 0 | 143 | 356 | 0 | 6 | 0 | 0 | 0 | 76 | 2 |

## UC

| Patients | Count | AS | LINC | Intronic | lncRNA | snRNA | tRNA | Divergent | miRNA | NE-MT | OT | rRNA | R60-Y | Small NF90 | snoRNA | Variant U1 |
|---|---|---|---|---|---|---|---|---|---|---|---|---|---|---|---|---|
| PC1 | 19126 | 558 | 935 | 24 | 171 | 3 | 0 | 122 | 274 | 0 | 6 | 0 | 0 | 0 | 66 | 0 |
| H1 | 16632 | 466 | 826 | 22 | 134 | 10 | 0 | 99 | 220 | 0 | 5 | 0 | 1 | 0 | 47 | 5 |
| H2 | 20692 | 600 | 1042 | 32 | 177 | 7 | 0 | 133 | 332 | 0 | 5 | 0 | 0 | 0 | 60 | 3 |
| PC2 | 18810 | 549 | 963 | 32 | 158 | 4 | 0 | 121 | 249 | 0 | 5 | 0 | 1 | 0 | 51 | 2 |
| PC3 | 4115 | 94 | 140 | 2 | 31 | 0 | 0 | 18 | 42 | 0 | 3 | 0 | 0 | 0 | 9 | 0 |
| H3 | 17102 | 482 | 842 | 20 | 139 | 0 | 0 | 97 | 221 | 0 | 4 | 0 | 1 | 0 | 47 | 0 |
| PC4 | 11249 | 293 | 470 | 12 | 89 | 1 | 0 | 51 | 124 | 0 | 2 | 0 | 0 | 0 | 25 | 1 |

**Figure s12.** Total RNA distributions across each method for all patients

**ExCy**

| Patients | Count | AS | LINC | Intronic | lncRNA | snRNA | tRNA | Divergent | miRNA | NE-MT | OT | rRNA | R60-Y | Small NF90 | snoRNA | Variant U1 |
|---|---|---|---|---|---|---|---|---|---|---|---|---|---|---|---|---|
| PC1 | 16477 | 183 | 134 | 25 | 70 | 5 | 0 | 0 | 295 | 0 | 1 | 0 | 1 | 0 | 66 | 1 |
| H1 | 17737 | 215 | 146 | 32 | 74 | 3 | 0 | 0 | 333 | 0 | 1 | 0 | 1 | 0 | 66 | 1 |
| H2 | 17934 | 220 | 147 | 28 | 73 | 4 | 0 | 0 | 375 | 0 | 1 | 0 | 1 | 0 | 73 | 0 |
| PC2 | 17888 | 224 | 146 | 32 | 78 | 5 | 0 | 0 | 362 | 0 | 1 | 0 | 2 | 0 | 60 | 2 |
| PC3 | 16647 | 184 | 133 | 30 | 67 | 3 | 0 | 0 | 244 | 0 | 1 | 0 | 0 | 0 | 56 | 0 |
| H3 | 16806 | 192 | 130 | 28 | 68 | 2 | 0 | 0 | 305 | 0 | 1 | 0 | 0 | 0 | 70 | 1 |
| PC4 | 16421 | 184 | 127 | 26 | 78 | 3 | 0 | 0 | 283 | 0 | 1 | 0 | 1 | 0 | 53 | 0 |

**ExoEasy**

| Patients | Count | AS | LINC | Intronic | lncRNA | snRNA | tRNA | Divergent | miRNA | NE-MT | OT | rRNA | R60-Y | Small NF90 | snoRNA | Variant U1 |
|---|---|---|---|---|---|---|---|---|---|---|---|---|---|---|---|---|
| PC1 | 16265 | 175 | 126 | 18 | 70 | 3 | 0 | 0 | 203 | 0 | 1 | 0 | 0 | 0 | 50 | 1 |
| H1 | 16902 | 188 | 137 | 26 | 71 | 3 | 0 | 0 | 221 | 0 | 1 | 0 | 0 | 0 | 60 | 1 |
| H2 | 15131 | 156 | 115 | 19 | 59 | 0 | 0 | 0 | 160 | 0 | 1 | 0 | 0 | 0 | 42 | 0 |
| PC2 | 15003 | 152 | 104 | 14 | 61 | 5 | 0 | 0 | 220 | 0 | 1 | 0 | 1 | 0 | 48 | 0 |
| PC3 | 14368 | 155 | 105 | 18 | 57 | 2 | 0 | 0 | 184 | 0 | 1 | 0 | 0 | 0 | 36 | 1 |
| H3 | 15980 | 177 | 120 | 22 | 61 | 1 | 0 | 0 | 217 | 0 | 1 | 0 | 0 | 0 | 51 | 0 |
| PC4 | 17002 | 197 | 136 | 26 | 67 | 1 | 0 | 0 | 296 | 0 | 1 | 0 | 1 | 0 | 73 | 1 |

**Fujifilm**

| Patients | Count | AS | LINC | Intronic | lncRNA | snRNA | tRNA | Divergent | miRNA | NE-MT | OT | rRNA | R60-Y | Small NF90 | snoRNA | Variant U1 |
|---|---|---|---|---|---|---|---|---|---|---|---|---|---|---|---|---|
| PC1 | 15494 | 187 | 117 | 17 | 66 | 4 | 0 | 0 | 232 | 0 | 1 | 0 | 0 | 0 | 51 | 0 |
| H1 | 18673 | 238 | 149 | 37 | 81 | 6 | 0 | 0 | 422 | 0 | 1 | 0 | 1 | 0 | 74 | 1 |
| H2 | 15838 | 171 | 119 | 26 | 69 | 4 | 0 | 0 | 258 | 0 | 1 | 0 | 0 | 0 | 46 | 1 |
| PC2 | 15186 | 158 | 115 | 20 | 63 | 2 | 0 | 0 | 201 | 0 | 1 | 0 | 0 | 0 | 48 | 0 |
| PC3 | 16438 | 194 | 125 | 23 | 70 | 4 | 0 | 0 | 253 | 0 | 1 | 0 | 2 | 0 | 62 | 0 |
| H3 | 15878 | 173 | 133 | 20 | 67 | 2 | 0 | 0 | 180 | 0 | 0 | 0 | 0 | 0 | 52 | 0 |
| PC4 | 17374 | 210 | 139 | 27 | 70 | 2 | 0 | 0 | 271 | 0 | 1 | 0 | 0 | 0 | 62 | 0 |

**UC**

| Patients | Count | AS | LINC | Intronic | lncRNA | snRNA | tRNA | Divergent | miRNA | NE-MT | OT | rRNA | R60-Y | Small NF90 | snoRNA | Variant U1 |
|---|---|---|---|---|---|---|---|---|---|---|---|---|---|---|---|---|
| PC1 | 14768 | 146 | 107 | 19 | 65 | 2 | 0 | 0 | 206 | 0 | 1 | 0 | 0 | 0 | 53 | 0 |
| H1 | 13046 | 132 | 96 | 17 | 56 | 5 | 0 | 0 | 167 | 0 | 1 | 0 | 1 | 0 | 36 | 0 |
| H2 | 15838 | 171 | 119 | 26 | 69 | 4 | 0 | 0 | 258 | 0 | 1 | 0 | 0 | 0 | 46 | 1 |
| PC2 | 14510 | 148 | 106 | 27 | 56 | 1 | 0 | 0 | 185 | 0 | 1 | 0 | 1 | 0 | 40 | 0 |
| PC3 | 3400 | 26 | 16 | 1 | 13 | 0 | 0 | 0 | 25 | 0 | 0 | 0 | 0 | 0 | 8 | 0 |
| H3 | 13417 | 129 | 98 | 17 | 54 | 0 | 0 | 0 | 159 | 0 | 1 | 0 | 1 | 0 | 36 | 0 |
| PC4 | 9097 | 77 | 62 | 10 | 37 | 0 | 0 | 0 | 84 | 0 | 0 | 0 | 0 | 0 | 18 | 0 |

**Figure s13.** Total RNA distributions across each method for all patients after mapping to Vesiclepedia.

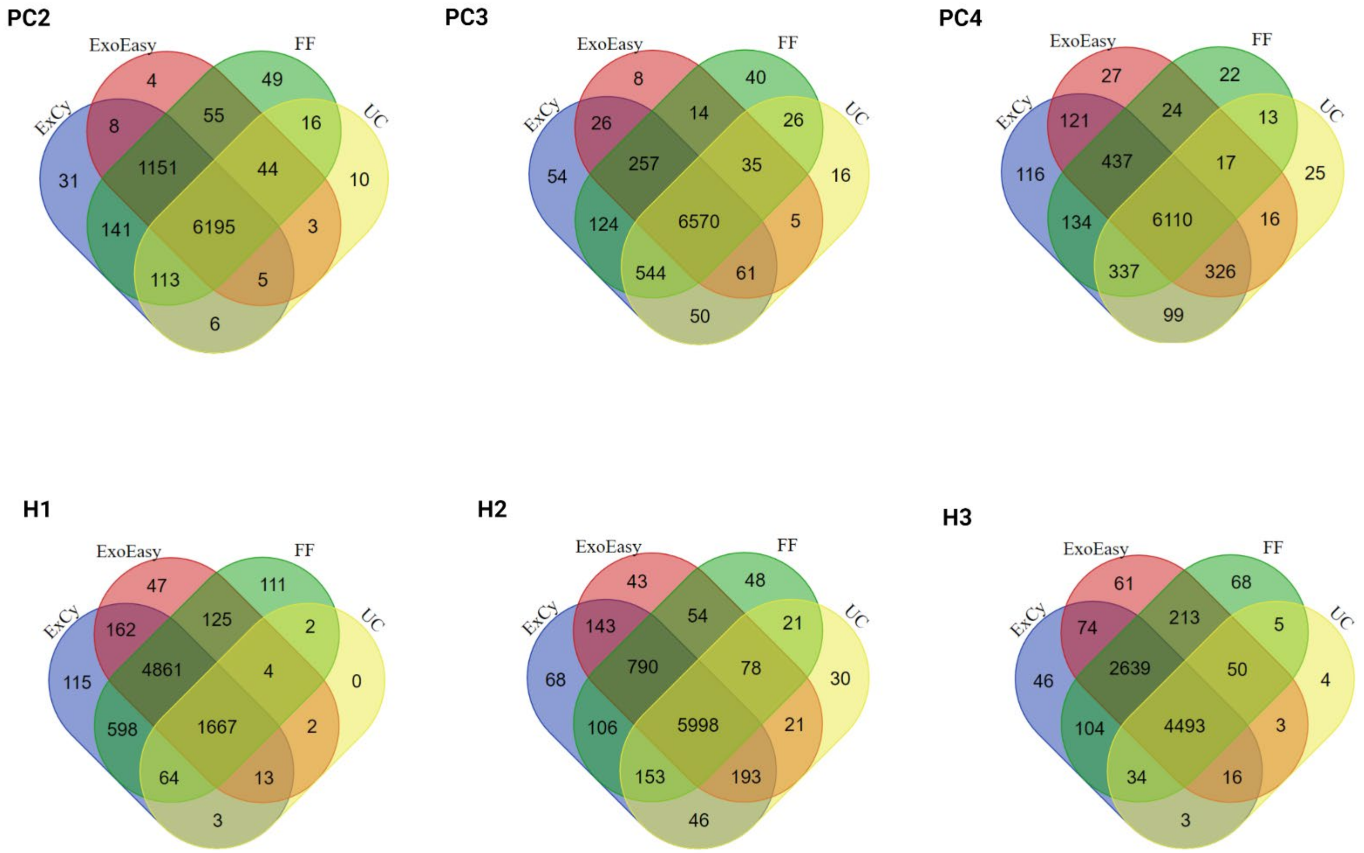


**Figure s14.** Venn diagrams indicating unique and shared mRNA transcripts across different methods for each patient.

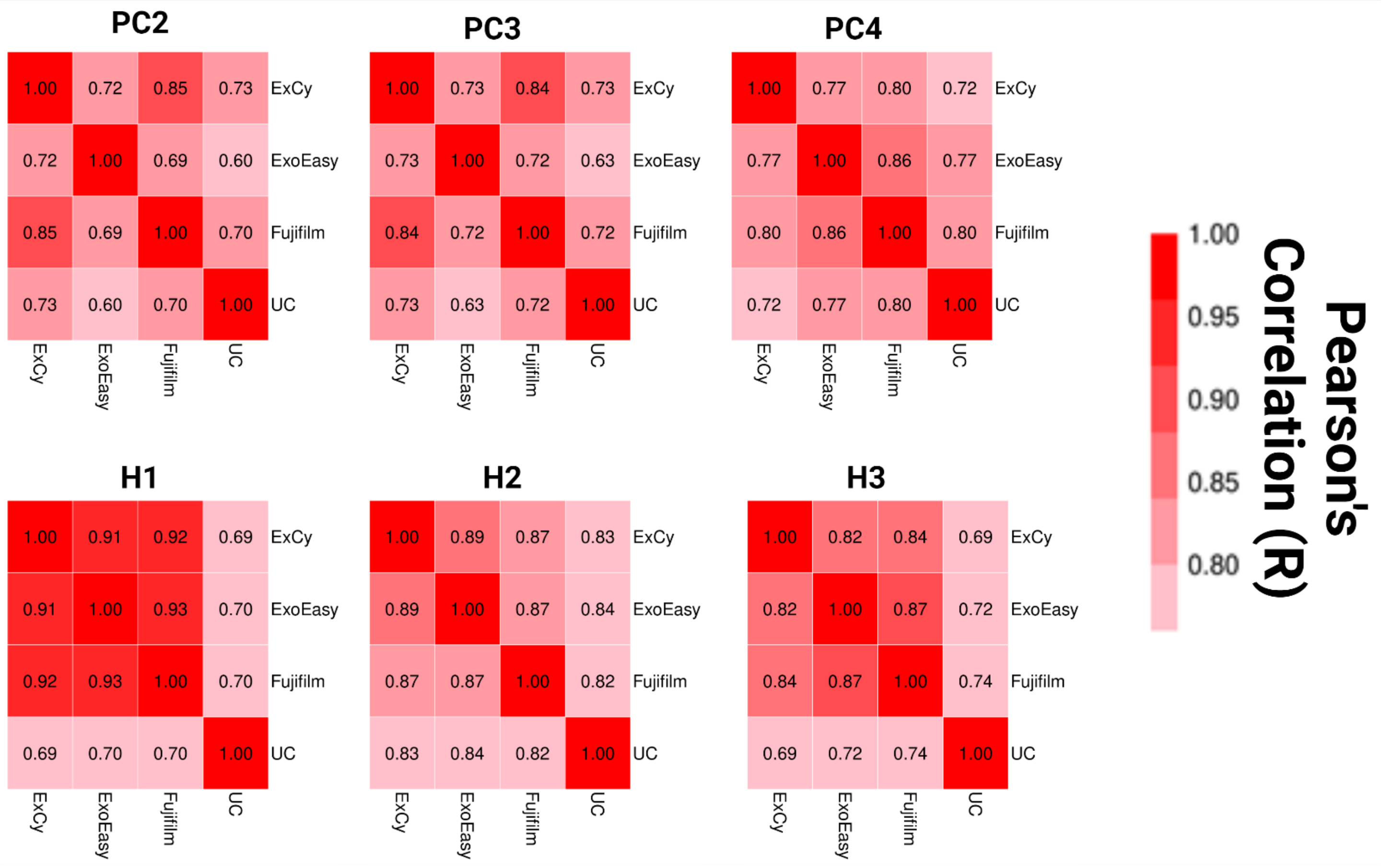


**Figure s15.** Pearson's correlation matrix across each method for all patients.

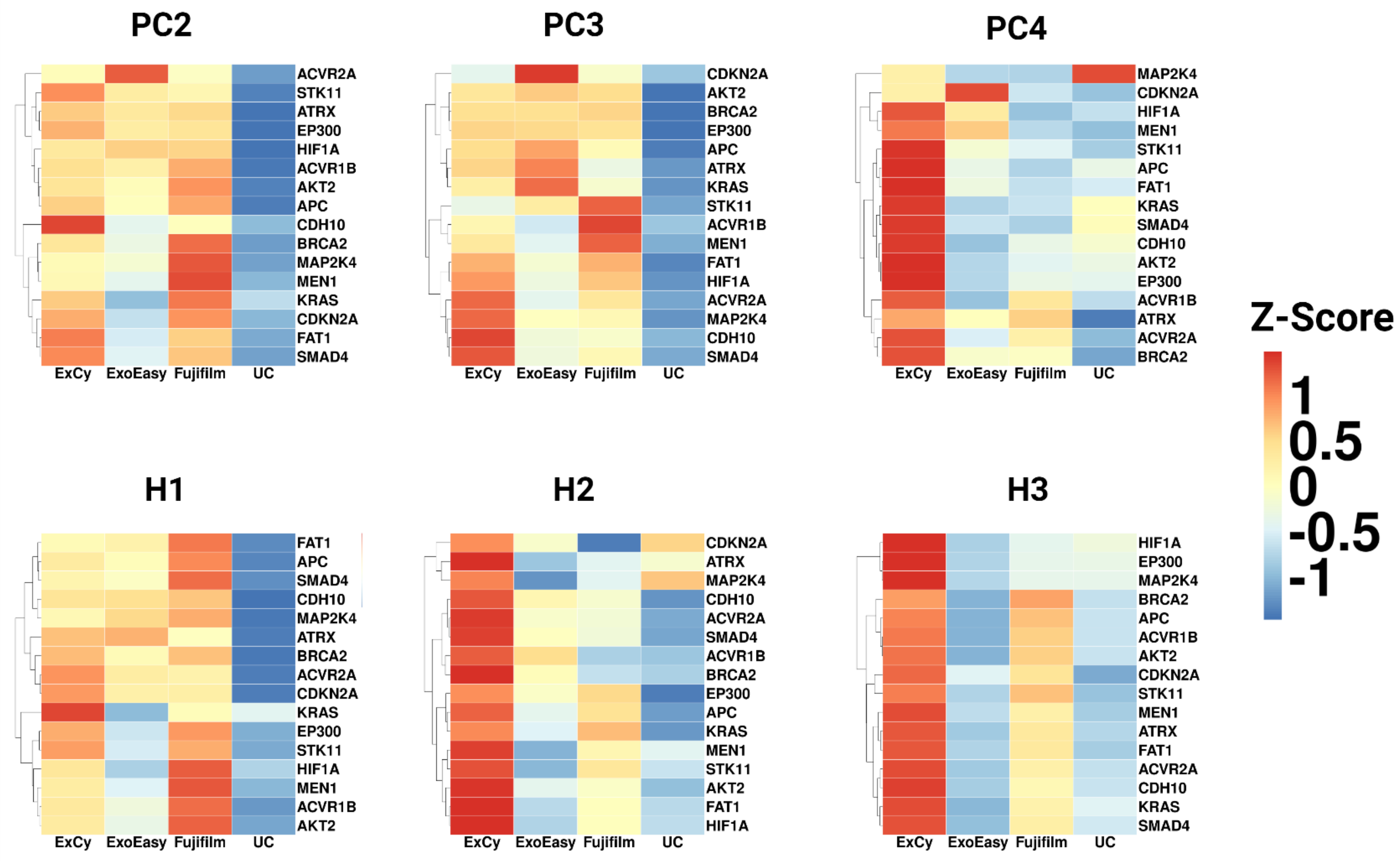


**Figure s16.** COSMIC pancreatic cancer heatmaps across all the other patients

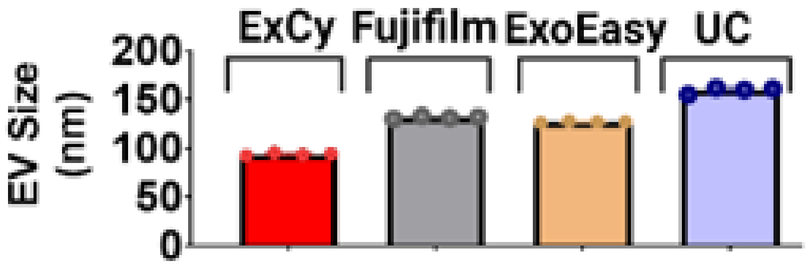


**Figure s17.** Summary of the EV sizes observed by transmission electron microscopy for figure 4.

| Transcripts | logFC | logCPM | LR | PValue | FDR |
|---|---|---|---|---|---|
| PUF60 | -9.2553 | 4.56867 | 32.16707 | 1.41E-08 | 0.00011 |
| VCX2 | -6.03026 | 2.1122 | 16.71289 | 4.35E-05 | 0.17194 |
| FTH1 | -3.21567 | 4.03465 | 14.19184 | 1.65E-04 | 0.3314 |
| PRSS8 | 1.827264 | 5.75381 | 14.16311 | 1.68E-04 | 0.3314 |
| CALML3 | 6.327308 | 1.45373 | 12.17029 | 4.86E-04 | 0.60367 |
| CTIF | 6.821779 | 2.27117 | 12.02635 | 5.25E-04 | 0.60367 |

**B) ExCy compared to Fujifilm**

| Transcripts | logFC | logCPM | LR | PValue | FDR |
|---|---|---|---|---|---|
| PUF60 | -9.4956 | 4.56867 | 33.50101 | 7.12E-09 | 5.63E-05 |
| VCX2 | -6.93387 | 2.1122 | 22.42687 | 2.18E-06 | 8.63E-03 |
| PAF1 | -1.69038 | 6.48951 | 12.17422 | 4.85E-04 | 7.14E-01 |
| CHORDC1 | -4.94693 | 1.58464 | 11.97977 | 5.38E-04 | 7.14E-01 |
| VCX | -4.38165 | 4.63627 | 11.78116 | 5.98E-04 | 7.14E-01 |

**C) ExCy compared to ExoEasy AND Fujifilm**

| Transcripts | logFC | logCPM | LR | PValue | FDR |
|---|---|---|---|---|---|
| PUF60 | -9.375448 | 4.568674 | 34.19844 | 4.98E-09 | **3.94E-05** |
| VCX2 | -6.482067 | 2.112198 | 21.0507 | 4.47E-06 | **1.77E-02** |
| SSX4 | 5.076693 | 1.790906 | 18.46378 | 1.73E-05 | **4.56E-02** |
| SAA1 | 7.032684 | 1.819769 | 17.71444 | 2.57E-05 | 5.07E-02 |
| CALML3 | 4.983599 | 1.453725 | 17.00773 | 3.72E-05 | 5.89E-02 |
| FTH1 | -2.894489 | 4.03465 | 13.35043 | 2.58E-04 | 3.41E-01 |

**D) Fujifilm compared to ExoEasy**

| Transcripts | logFC | logCPM | LR | PValue | FDR |
|---|---|---|---|---|---|
| SCG5 | 6.023166 | 1.53196 | 13.57541 | 0.00022917 | 1 |
| CTIF | -6.3206 | 2.27117 | 10.36676 | 0.00128305 | 1 |
| BAG6 | -7.18689 | 2.55276 | 9.770019 | 0.00177381 | 1 |
| DLST | -1.15268 | 7.08337 | 9.673911 | 0.00186903 | 1 |
| CYBA | 1.054164 | 6.89675 | 7.781736 | 0.0052777 | 1 |
| PDZRN4 | 0.960956 | 6.81347 | 7.279409 | 0.00697495 | 1 |

**Figure s18.** Investigating the differential analysis of EVs isolated by ExCy, ExoEasy, and Fujifilm from healthy samples to understand marker discovery differences with edgeR. **A)** ExCy was compared to ExoEasy and **B)** Fujifilm separately, then **C)** compared to the combined statistical effect of ExoEasy and Fujifilm. **D)** Fujifilm compared to ExoEasy.

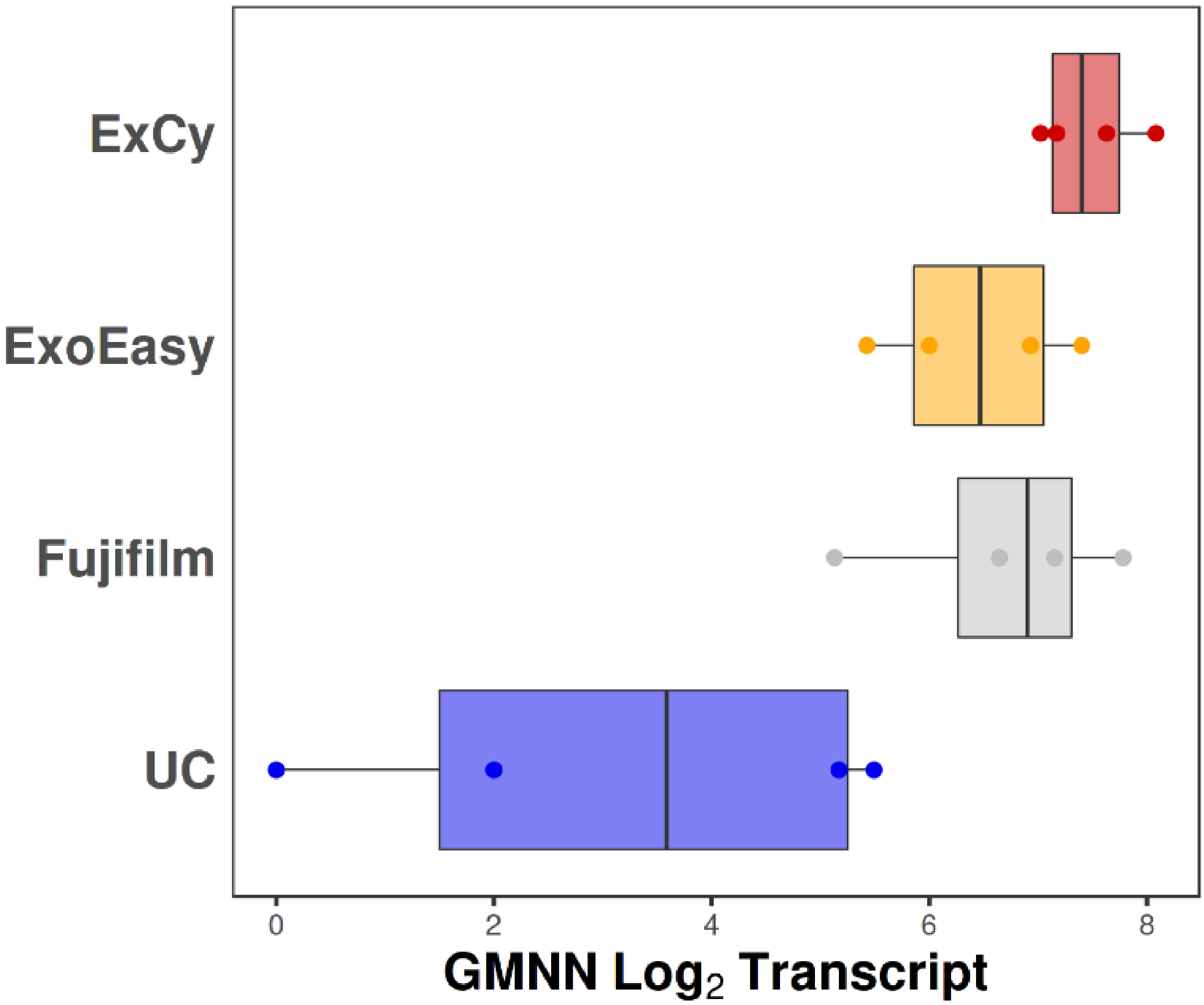


**Figure s19.** GMNN mRNA levels across the four EV isolation methods to understand interaction with PHC1. If PHC1 was detected in an EV, then GMNN must likely be isolated along with PHC1, since GMNN is regulated by PHC1 in the pancreas.
